# Global dynamics of a general vector-borne disease model with two transmission routes

**DOI:** 10.1101/486720

**Authors:** Sk Shahid Nadim, Indrajit Ghosh, Joydev Chattopadhyay

## Abstract

In this paper, we study the dynamics of a vector-borne disease model with two transmission paths: direct transmission through contact and indirect transmission through vector. The direct transmission is considered to be a non-monotone incidence function to describe the psychological effect of some severe diseases among the population when the number of infected hosts is large and/or the disease possesses high case fatality rate. The system has a disease-free equilibrium which is locally asymptomatically stable when the basic reproduction number (*R*_0_) is less than unity and may have up to four endemic equilibria. Analytical expression representing the epidemic growth rate is obtained for the system. Sensitivity of the two transmission pathways were compared with respect to the epidemic growth rate. We numerically find that the direct transmission coefficient is more sensitive than the indirect transmission coefficient with respect to *R*_0_ and the epidemic growth rate. Local stability of endemic equilibria is studied. Further, the global asymptotic stability of the endemic equilibrium is proved using Li and Muldowney geometric approach. The explicit condition for which the system undergoes backward bifurcation is obtained. The basic model also exhibits the hysteresis phenomenon which implies diseases will persist even when *R*_0_ < 1 although the system undergoes a forward bifurcation and this phenomenon is rarely observed in disease models. Consequently, our analysis suggests that the diseases with multiple transmission routes exhibit bi-stable dynamics. However, efficient application of temporary control in bi-stable regions will curb the disease to lower endemicity. In addition, increase in transmission heterogeneity will increase the chance of disease eradication.

## 1. Introduction

The mode of transmission of some infectious diseases may be through both direct and indirect contact. In direct transmission, a pathogen can infect a susceptible host through infected host, and in indirect transmission, a pathogen can infect a susceptible host through an infected vector. For instance, Middle East respiratory syndrome coronavirus (MERS-CoV) (1) and Zika virus (2) may be spread both directly by person-to-person contact and indirectly through a vector. Understanding the transmission dynamics of such diseases is of utmost interest. Mathematical models of epidemic diseases have been very insightful in uncovering epidemiology and transmission patterns of the diseases. It is very important to understand such epidemic patterns for better public health interventions and control of outbreaks. Another important objective of these models is to inform public health organizations about the effect of control policies on a certain population. However, hosts behaviour and psychological factors are sometimes the main drivers of further transmission of the disease. For example, the epidemic outbreak of severe acute respiratory syndrome (SARS) (3; 4) had such psychological effects and individual protection policies have been found very effective (5; 6) in decreasing the rate of infection at the late stage of SARS outbreak, even when the number of infective individuals were getting larger relatively. Additionally, the influenza pandemic of 2009 has triggered a psychological effect that caused adoption of preventive measures by a significant portion of the population (4). Recent MERS-CoV outbreak in Arabian peninsula also imposed a significant behaviour change in the local people (1). Thus, psychological effects are necessary in modelling infectious disease transmission. The general incidence rate in this context was proposed by Lui et al (7) of the form 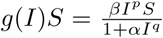, where *βI*^*p*^ represent the force of infection of the diseases, 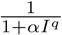 represent the psychological effect from the behavioral changes of susceptible population when the number of infected populations increases, *p, q*, and *β* are positive constant and *α* is a nonnegative constant. Ruan and Wang (8) studied an SIRS epidemic model with a specific nonlinear incidence rate 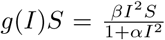. Afterwards, Xiao and Ruan (9) studied the dynamical behaviour of an SIR epidemic model with the incidence function 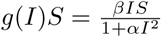.

However, the dynamics of an epidemic model with two transmission routes where the direct transmission incorporates psychological effect has not been studied so far. The ongoing MERS-CoV epidemic and zika virus epidemic are examples of diseases with two routes of transmission having psychological effects. The psychosocial implication on the disease outbreak is more dangerous than any other physical effect. West Nile Virus outbreak in the US is a good example for the onset of psychogenic illness (10) and may generate mistrust towards national or health authorities. For example, the mild effects of Zika virus, there is a correlation between psychosis with fetus microcephaly where pregnant women and their relatives are highly sensitive group. Therefore, there is a negative correlation between the psychological effect and disease prevalence and hence we consider non-monotone incidence function i.e *p* < *q* to describe the direct transmission between hosts in our study. This type of function can be used to interpret the “psychological effects” (11). We use the incidence rate 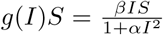 where *βI* represent the force of infection of the diseases, 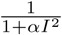 represent the psychological effect from the behavioral changes of susceptible population when the number of infected populations increases. Our goal is to develop a simple mathematical model that capture a range of disease dynamics with multiple mode of transmissions. Moreover, we would like to perform a detailed stability and bifurcation analysis of the proposed model. By using analytical methods we intend to uncover the dynamical properties of the diseases with two routes of transmission.

The rest of the paper is organized as follows: Section 2 consist of the model formulation and the basic assumptions. In Section 3, preliminary mathematical analysis of the proposed model is carried out. Section 4 is devoted to the global stability of the endemic equilibrium using geometric approach while in Section 5 backward bifurcation and forward bifurcation with hysteresis effect is presented. In Section 6 we investigate some numerical scenarios such as effect of temporary control on bistable dynamics, effect of the strength of psychological fear and impact of heterogeneity in indirect transmission. Finally, discussion and conclusion is presented in Section 7.

## 2. Model formulation

We consider two population groups as host and vector population. Time-dependent state variables are taken to describe the compartments of the two populations. Let at time t, *N*_*h*_(*t*) represents total host population and *N*_*v*_(*t*) represents total vector population. We consider a ‘SIR’ type model for the host population and a ‘SI’ type model for the vectors. The host population is subdivided into three mutually disjoint classes: susceptible, *S*_*h*_; infected, *I*_*h*_; recovered, *R*_*h*_. Thus, at any time t, the size of the host population is given by *N*_*h*_ = *S*_*h*_ + *I*_*h*_ + *R*_*h*_. The vector population consist of two classes namely, susceptible vectors, *S*_*v*_; and infected vectors, *I*_*v*_. With this division, the size of the vector population at time *t* is given by *N*_*v*_(*t*) = *S*_*v*_(*t*) + *I*_*v*_(*t*). We assume all newborn hosts are fully susceptible. The susceptible host population increases at a constant rate Π_*h*_. The susceptible population decreases due to getting infection from infected vectors and infected hosts and natural mortality at a rate *µ*_*h*_. Therefore the average life time of host population is 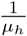. The model incorporates a non-monotone incidence function for the direct transmission. The force of infection in this regard is considered as 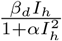, where *β*_*d*_ and *α* are defined in Table 1. The infected host population *I*_*h*_ decreases as a result of death and recovery. It is assumed that the recovered hosts get a life-long immunity. *γ* is the recovery rate of the infected hosts. Recovered host are decreased at a rate *µ*_*h*_. The susceptible vector population increases at a constant rate Π_*v*_ and decreases by getting infection from infected host and by natural mortality of vectors. Standard forces of infection is taken between host and vector populations, respectively, 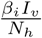 and 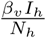. The parameter *β*_*i*_ is the effective transmission coefficient between susceptible host population and infected vector population, and the parameter *β*_*v*_ is the effective transmission coefficient of the infected host population and susceptible vector population. Taking the above considerations into account, the governing equations of the disease dynamics is given by:

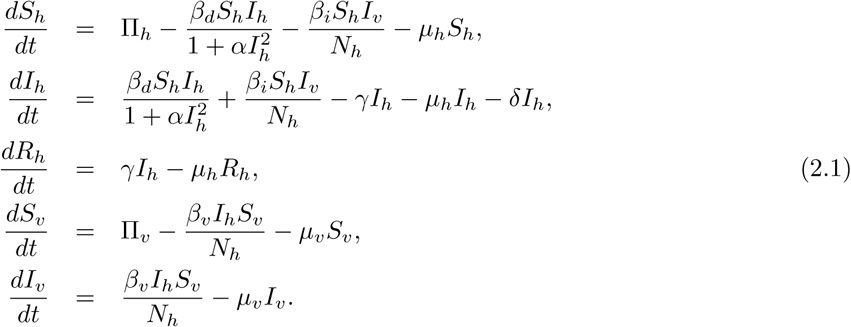

**Table 1:** Description of parameters used in the model

| Parameters | Interpretation |
| --- | --- |
| $\Pi_h$ | Recruitment rate of hosts |
| $\Pi_v$ | Recruitment rate of vectors |
| $\mu_h$ | Natural host death rate |
| $\mu_v$ | Natural vector death rate |
| $\beta_d$ | Transmission coefficient from infected host to susceptible host |
| $\beta_i$ | Transmission coefficient from infected vector to susceptible host |
| $\beta_v$ | Transmission coefficient from infected host to susceptible vector |
| $\alpha$ | Strength of psychological effect |
| $\gamma$ | Recovery rate of infected host |
| $\delta$ | Disease induced mortality rate |

Interpretations of all the model parameters and their biological meanings are given in the Table 1.

## 3. Mathematical analysis

### 3.1. Basic properties of the model

We assume that the vector and host state variables are non-negative for all *t* ≥ 0. Let

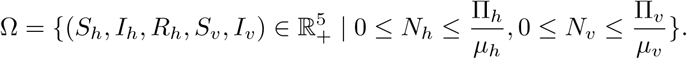

We claim the following result.

#### Theorem 3.1.

Whenever the initial conditions are non-negative, model (2.1) has non-negative solutions and the region 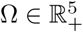 is positively invariant and globally attracting for the above system.

#### Proof.

We can rewrite the system (2.1) in the following form

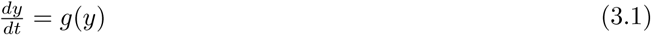

with *y* = (*y*_1_, *y*_2_, *y*_3_, *y*_4_, *y*_5_) = (*S*_*h*_, *I*_*h*_, *R*_*h*_, *S*_*v*_, *I*_*v*_) and *g*(*y*) = (*g*_1_(*y*), *g*_2_(*y*), *g*_3_(*y*), *g*_4_(*y*), *g*_5_(*y*)) denote the right hand side of the functions as in the following

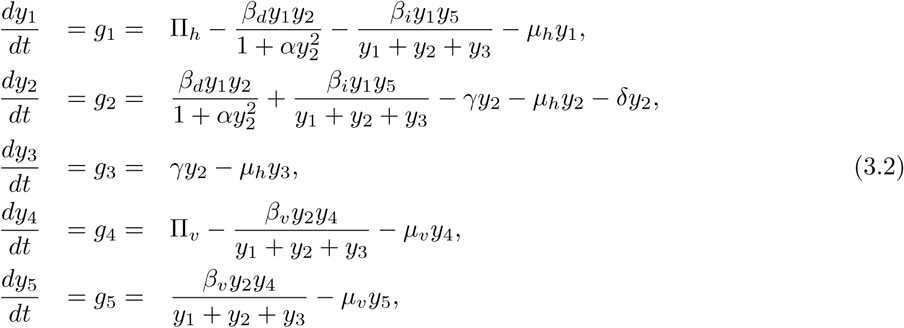

Clearly, for every *j* = 1, *…,* 5, *g*_*j*_(*x*) *≥* 0 if 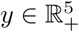 and *y*_*j*_ = 0. Since the host and vector populations are non-negative, thus the right hand side of the system (3.1) is locally Lipschitz in Ω. Following (12) and (13), we see that the system (2.1) has a unique solution. The following equations are satisfied by the total host and vector populations,

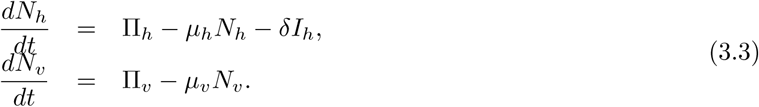

Since 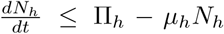, it implies that 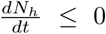 if 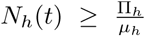 and 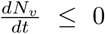 if 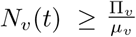. Consequently, it can be shown that 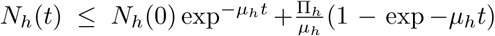 and 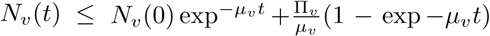. In particular, 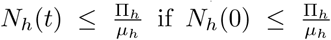 and 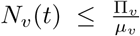 if 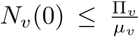. Thus the region Ω is positively invariant. Further, if 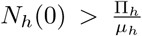 and 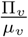, then the solutions of the system (2.1) either enter Ω in finite time or *N*_*h*_(*t*) approaches 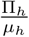 and *N*_*v*_(*t*) approaches 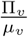 asymptomatically. Hence, all solutions in 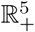 will eventually enter Ω.

### 3.2. Basic reproduction number and disease free equilibrium

The basic reproduction number, *R*_0_, is the number of secondary infections which one infected individual would produce in a entirely susceptible population. The disease free equilibrium (DFE) of the system is given by 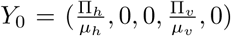. There are two infected compartments in the model (2.1), *I*_*h*_ and *I*_*v*_. *R*_0_ can be derived from the jacobian matrix of the system (2.1) calculated at *Y*_0_ together with the assumption of local asymptotic stability of *Y*_0_ and is given by

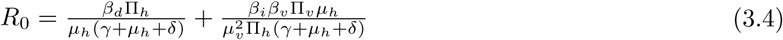

#### Theorem 3.2.

The DFE Y_0_ of the system (2.1) is locally asymptotically stable, if R_0_ < 1, and unstable if R_0_ > 1.

#### Proof.

The Jacobian of the system (2.1) at DFE *Y*_0_ is given as follows

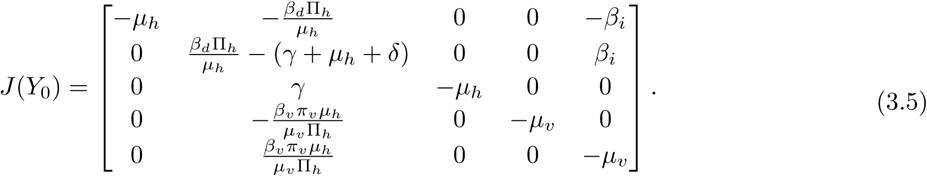

Then the characteristic equation of (3.5) is given by

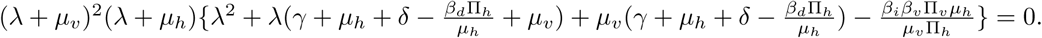

The eigenvalues of this jacobian matrix is

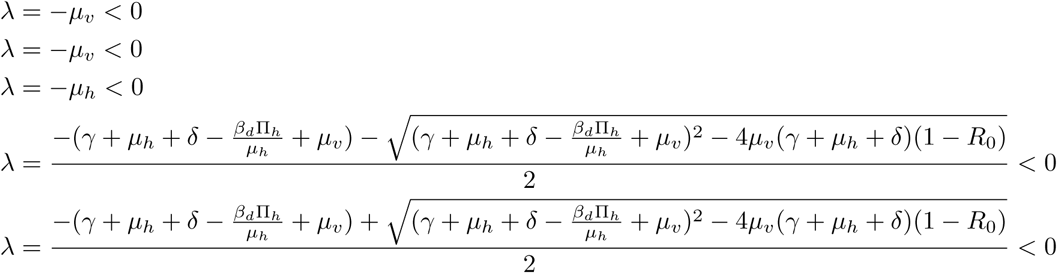

for *R*_0_ < 1. Hence the theorem is proved.

It is well established that the threshold *R*_0_ governs the local stability of DFE. For *R*_0_ < 1 DFE is locally asymptotically stable and it is unstable when *R*_0_ > 1 (14; 15). Further, when *R*_0_ > 1, the disease is persistent. The epidemic growth rate (EGR) remains a useful quantity for estimating the transmission potential at the population level. The initial EGR is defined as the dominant eigenvalue of the Jacobian at DFE. In the following subsection we calculate the EGR for the system 2.1. This will let us infer about the contribution of different transmission pathways on the epidemic growth rate, and will give useful results for calculating *R*_0_ from an observed epidemic growth rate.

### 3.3. Epidemic growth rate

Following Tien and Earn (14), we calculate the initial EGR for the system (2.1). From this we can infer about the contribution of the two transmission routes with respect to the EGR.

Let *λ* be the initial EGR for the model (2.1). When *R*_0_ > 1, the dominant eigenvalue of the Jacobian at the DFE for model (2.1) is given by

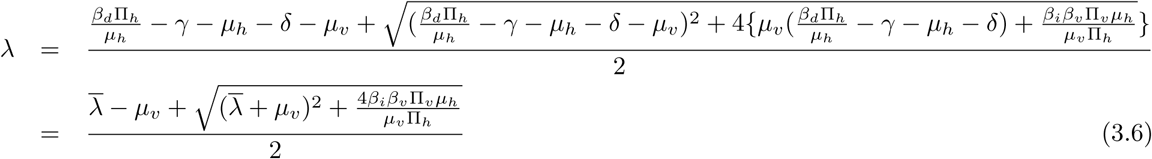

where

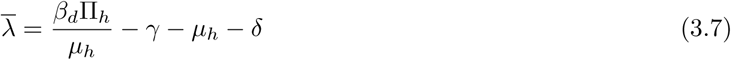

Note that 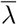 is the EGR for the ‘SIR’ sub-model of the system (2.1) and is given by

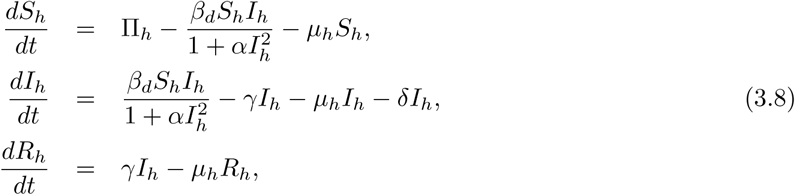

If there is no indirect transmission through vector (*β*_*i*_ → 0), then the system (2.1) becomes the sub-model (3.8) and 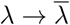. Additionally, from (3.6), it can be noted that

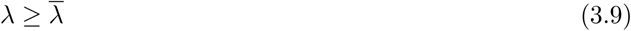

The mean time spent in the infected host class is dependent on the host lifetime, the mean infectious period and the disease induced death rate. Similarly, the mean vector lifetime is is the mean time spent in the infected vector class. Therefore, let 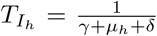 denotes the mean time spent in the infected class, and let 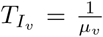 denotes the mean vector lifetime. Rearranging the equation (3.6), we get the following expression interlinking *R*_0_ and *λ*:

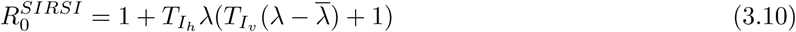

We observe that equation (3.10) involves both 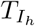 and 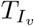. If there is no vector transmission, this equation converges to the relationship between *R*_0_ and EGR for the SIR model (3.8):

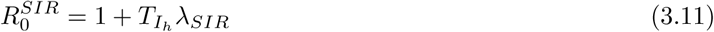

where 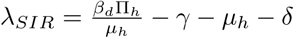

Empirical estimate of *R*_0_ from a given disease outbreak data is of considerable interest in quantitative epidemiology (16). Assume that the observed EGR (*λ*_*obs*_) is available, and let the parameters be chosen for the ‘SIR’ and ‘SIRSI’ systems such that the EGR calculated from the model and the observed EGR *λ*_*obs*_ is same for these two models. Let us denote the value for *R*_0_ in ‘SIRSI’ system by 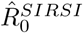 and let 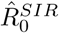 denotes the value for *R*_0_ in ‘SIR’ system. Then we claim the following result which describes that when the parameters *γ, µ*_*h*_ and *δ* are known, 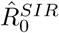 value is less than or equal to 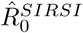.

#### Proposition 1.

Let λ_obs_ > 0 is known. Now we consider the suitable parameters for the ‘SIR’ and ‘SIRSI’ systems, such that the EGRs are equal to λ_obs_ for both models. Let 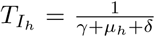 be the equal for both the models. Then 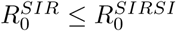.

#### Proof.

From (3.10), we have 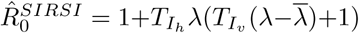. From (3.11), we have 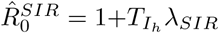.

Subtracting these two expressions gives:

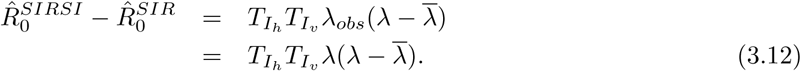

Provided, the observed growth rate *λ*_*obs*_ is equal to the EGR *λ* for the ‘SIRSI’ model (2.1). Since 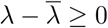 (3.9), then 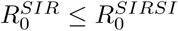 as claimed.

### 3.4. Comparing the two transmission routes

We are now interested in finding the relative contributions of the person-person and vector-person transmission pathways on disease dynamics. To investigate this, we can vary *β*_*d*_ and *β*_*i*_ keeping *R*_0_ fixed (14). For *R*_0_ > 1, the following result shows that the rate of change of EGR with respect to *β*_*i*_ is negative.

#### Proposition 2.

Let R_0_ > 1 and R_0_ is fixed for system (2.1). Let 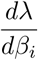 denote the total derivative of λ with respect to β_i_. Then 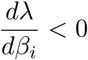.

#### Proof.

The EGR *λ* is explicitly dependent on both *β*_*i*_ and *β*_*d*_. From the analytical expression of *R*_0_, *β*_*d*_ can be determined in terms of *β*_*i*_ as, 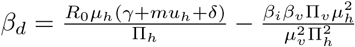. The term 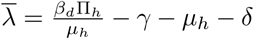 (3.7) is involved in the expression of 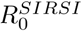 given by (3.10). We will calculate the total derivative of 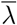:

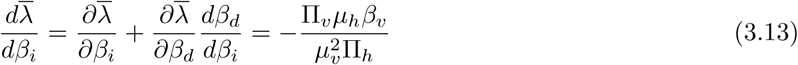

Since *R*_0_ is fixed and using (3.13), the total derivative of (3.10) gives

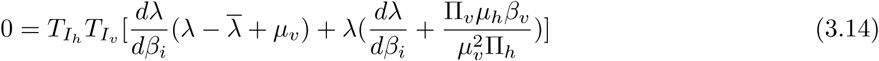

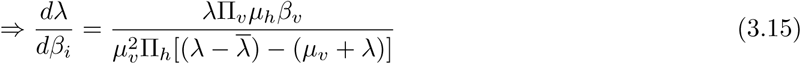

We have 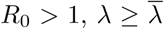 and *λ* > 0. Finally, *µ*_*v*_ > 0. Thus, the numerator become positive and the denominator is negative in (3.15), giving that 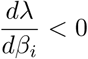.

Now we calculate the relationship of 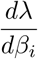 with 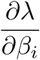 and 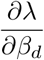 as follows:

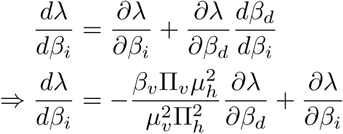

#### Corollary 3.1.

Let R_0_ > 1 and the initial EGR λ be given by (3.6). If 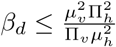, then 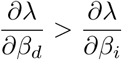.

To verify this result numerically, sensitivity index of *λ* with respect to the parameters *β*_*d*_ and *β*_*i*_ are computed. The normalized forward sensitivity index of a quantity with respect to a parameter is defined as the ratio of the relative change in the variable to the relative change in the parameter. Mathematically, the normalized forward sensitivity index of a variable *m* that depends explicitly on a parameter *τ* is defined as:

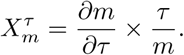

The normalized forward sensitivity indices of *λ* with respect to the parameters *β*_*d*_ and *β*_*i*_ are found to be

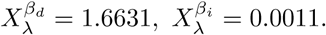

for the parameters value Π_*h*_ = 5, *β*_*d*_ = 0.1, *β*_*v*_ = 0.1, *µ*_*h*_ = 0.1, *µ*_*v*_ = 0.5, Π_*v*_ = 30, *d* = 1, *γ* = 0.9, *α* = 1, *β*_*i*_ = 0.1.

The fact that 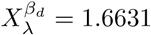 means that 1% increase in *β*_*d*_, keeping other parameters fixed, will produce 1.6631% increase in *λ*. On the other hand, increasing the parameter *β*_*i*_ by 1%, keeping the other parameters constant, the value of *λ* increases by 0.0011%. Additionally, we computed the normalized sensitivity indices of *R*_0_ with respect to the parameters *β*_*d*_ and *β*_*i*_. In this case, increasing the parameters *β*_*d*_ and *β*_*i*_ by 1%, the value of *R*_0_ increases by 0.9952% and 0.0048;% respectively. These results suggest that the direct transmission pathway is more sensitive to EGR and *R*_0_ than the indirect one, which indicates that the direct transmission control will be more effective in halting the early phase of the epidemic.

## 4. Endemic equilibria and its stability

In this section we discuss the existence of endemic equilibria and its stability conditions. Let 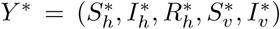 be an arbitrary endemic equilibrium of the system(2.1). Therefore, equating the right hand sides of the equations of system (2.1) to zero, we have

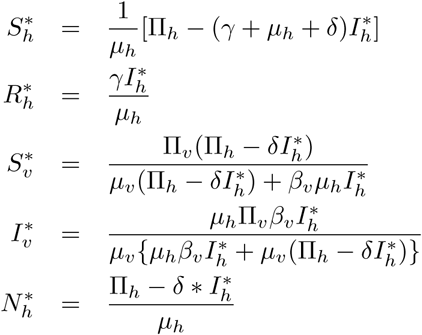

and 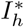 comes from the equation

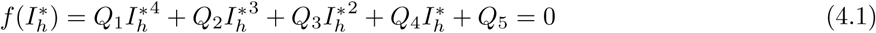

where,

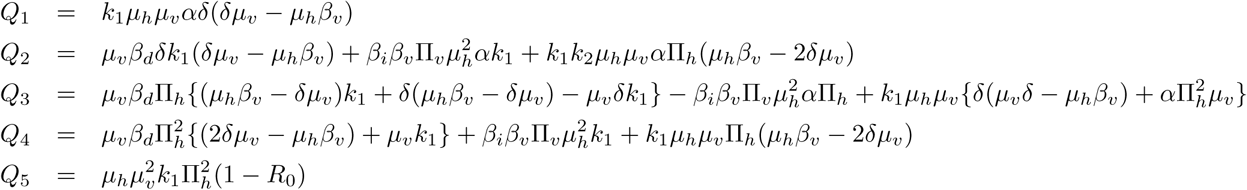

Here

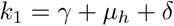

Since all parameters are positive and if *R*_0_ < 1, then *Q*_5_ > 0. Hence, depending on the signs of *Q*_1_, *Q*_2_, *Q*_3_ and *Q*_4_, the polynomial(4.1) can have at most four positive real roots. We use Descartes rule of signs to enlist the possible number of positive real roots of the equation 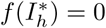; Table 2 and Table 3.

**Table 2:** Various possibilities for positive real roots of 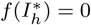 for *R*_0_ > 1

| Case | $Q_1$ | $Q_2$ | $Q_3$ | $Q_4$ | $Q_5$ | # sign changes | # possible positive real roots |
| --- | --- | --- | --- | --- | --- | --- | --- |
| 1 | + | + | + | + | - | 1 | 1 |
| 2 | + | + | + | - | - | 1 | 1 |
| 3 | + | + | - | + | - | 3 | 1,3 |
| 4 | + | - | + | + | - | 3 | 1,3 |
| 5 | - | + | + | + | - | 2 | 0,2 |
| 6 | + | + | - | - | - | 1 | 1 |
| 7 | + | - | - | + | - | 3 | 1,3 |
| 8 | + | - | + | - | - | 3 | 1,3 |
| 9 | - | - | + | + | - | 2 | 0,2 |
| 10 | + | - | - | - | - | 1 | 1 |
| 11 | - | + | - | - | - | 2 | 0,2 |
| 12 | - | - | + | - | - | 2 | 0,2 |
| 13 | - | - | - | + | - | 2 | 0,2 |
| 14 | - | - | - | - | - | 0 | 0 |
| 15 | - | + | - | + | - | 4 | 0,2,4 |
| 16 | - | + | + | - | - | 2 | 0,2 |

**Table 3:** Various possibilities for positive real roots of 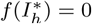 for *R*_0_ < 1

| Case | $Q_1$ | $Q_2$ | $Q_3$ | $Q_4$ | $Q_5$ | # sign changes | # possible positive real roots |
| --- | --- | --- | --- | --- | --- | --- | --- |
| 1 | + | + | + | + | + | 0 | 0 |
| 2 | + | + | + | - | + | 2 | 0,2 |
| 3 | - | - | - | + | + | 1 | 1 |
| 4 | - | - | - | - | + | 1 | 1 |
| 5 | - | - | + | + | + | 1 | 1 |
| 6 | + | - | - | - | + | 2 | 0,2 |
| 7 | - | + | - | + | + | 3 | 1,3 |
| 8 | - | + | - | - | + | 3 | 1,3 |
| 9 | - | - | + | - | + | 3 | 1,3 |
| 10 | + | + | - | + | + | 2 | 0,2 |
| 11 | + | + | - | - | + | 2 | 0,2 |
| 12 | + | - | + | + | + | 2 | 0,2 |
| 13 | + | - | + | - | + | 4 | 0,2,4 |
| 14 | - | + | + | + | + | 1 | 1 |
| 15 | - | + | + | - | + | 3 | 1,3 |
| 16 | + | - | - | + | + | 2 | 0,2 |

### Theorem 4.1.

The system (2.1) has unique endemic equilibrium Y ^*^ if R_0_ > 1 and cases 1, 2, 6 and 10 in Table 2 are satisfied, and has unique endemic equilibrium Y ^*^ if R_0_ < 1 and cases 3, 4, 5 and 14 in Table 3 are satisfied. The system (2.1) could have more than one endemic equilibrium if R_0_ > 1 and cases 3 --5, 7 --9, 11 --13, 15 and 16 in Table 2 are satisfied, and could have more than one endemic equilibrium if R_0_ < 1 and cases 2, 6 --13, 15 and 16 in Table 3 are satisfied.

### 4.1. local stability of endemic equilibrium

The Jacobian matrix of the system (2.1) evaluated at endemic equilibrium *Y* ^*\**^ can be written as

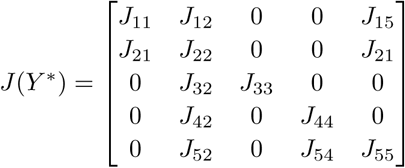

where

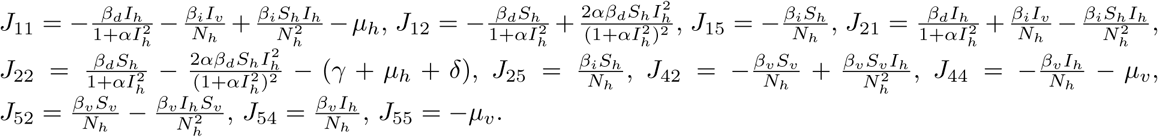

Clearly, *-µ*_*h*_ is one root of *J* (*Y* ^*\**^), which is negative. The remaining roots can be determined from the following characteristic equation which is given by

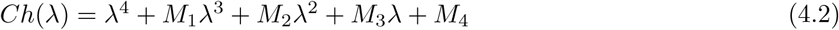

where

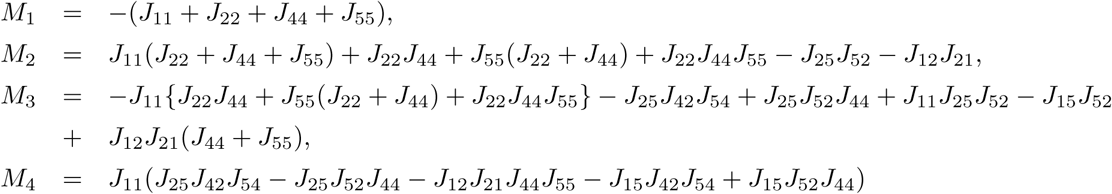

Using the RouthHurwitz criterion, the roots of equation (4.2) are either negative or have negative real parts if and only if following conditions are satisfied

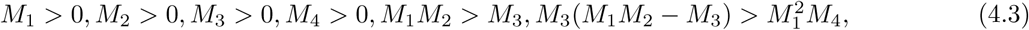

Thus, the endemic equilibrium *Y*^*\**^ is locally asymptotically stable if and only if conditions in (4.3) are satisfied.

### 4.2. Global stability of endemic equilibrium

In Theorem 4.1, there exist conditions for which the system (2.1) possesses unique endemic equilibrium. We will use the geometric approach proposed by Li and Muldowney to study the global stability of the endemic equilibrium (17; 18; 19). For this it is necessary to reduce the dimension of the system (2.1) by at least one as this method is applicable to systems whose order is at most four. Since 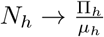 as *t* → *∞* (20), the system (2.1) reduces to the limit system

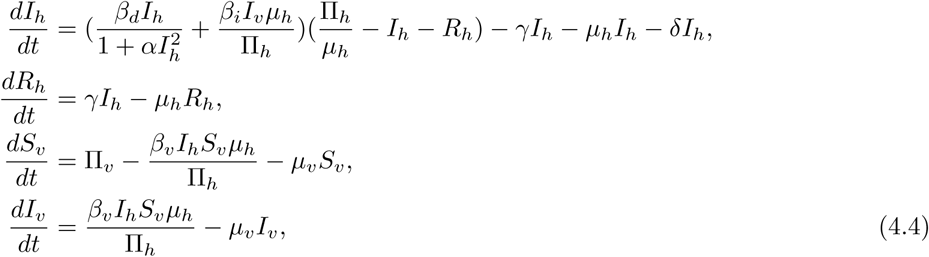

Clearly, the solutions of the limit system (4.4) with non-negative initial conditions remain non-negative. Therefore, We can study the model in the following region

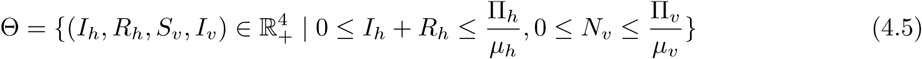

However, before going to the main result, let us discuss some preliminary results and definitions. Consider the autonomous dynamical system:

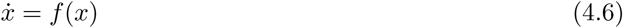

where *f*: *D* → *R*^*n*^, *D ⊂ R*^*n*^ open set and simply connected and *f ∈ C*^1^(*D*). Let *x*^*\**^ be the solution of (4.5) such that *x*(0, *x*_0_) = *x*_0_, i.e, *f* (*x*^*\**^) = 0. We recall that *x*^*\**^ is said to be globally stable in *D* if it is locally stable and all trajectories in *D* converges to *x*^*\**^.

#### Definition

The set *K* is absorbing subset in *D* for the system (4.5), if for every compact *K*_1_ *⊂ D, x*(*t, K*_1_) *⊂ K* for sufficiently large *t*, where

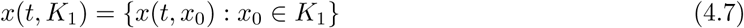

Let *Q*(*x*) be a matrix of size 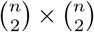 and *Q ∈ C*^1^(*D*). Suppose that *Q*^*-*1^ exists and it is continuous in *K.* Where *K* is a compact absorbing set in *D*.

We define

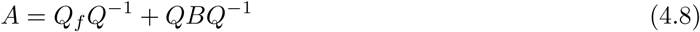

where *Q*_*f*_ is obtained replacing every entry *q*_*ij*_ of *Q* by its directional derivatives with respect the vectorial field *f*.

The Lozinskii measure *µ*(*A*) with respect to the norm ||.|| in *R*^*N*^, 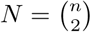, is defined as (18),

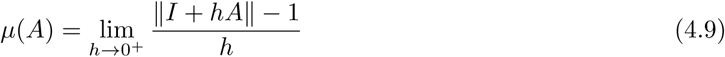

We will apply the following (18)

#### Theorem 4.2.

If K is a compact absorbing subset in the interior of D, and there exist ν > 0 such that the Lozinskii measure µ(A) ≤ –ν for all x ∈ K, Then every omega limit point of system (4.4) in the interior of D is an equilibrium in K.

The system (2.1) admits exactly one endemic equilibrium *Y* ^*\**^ when *R*_0_ > 1 for the cases 1, 2, 6, 10 in Table 2. Further, we know that the DFE *Y*_0_ is unstable when *R*_0_ > 1. The instability of *P*_0_, together with *P*_0_ *∈ ∂*Θ, which implies the uniform persistence of the state variable (21). Thus there exists a constant *c* > 0 such that any solution (*I*_*h*_(*t*), *R*_*h*_(*t*), *S*_*v*_(*t*), *I*_*v*_(*t*)) with (*I*_*h*_(0), *R*_*h*_(0), *S*_*v*_(0), *I*_*v*_(0)) in the orbit of system (4.4) satisfies

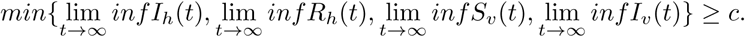

The uniform persistence of the system (4.4), incorporating the boundedness of Θ, suggests that the compact absorbing set in the interior of Θ;see (22). Hence the above Theorem 4.2 may be applied with *D* = Θ.

#### Theorem 4.3.

Let R_0_ > 1, the conditions 1,2,6,10 in Table 2 are satisfied and 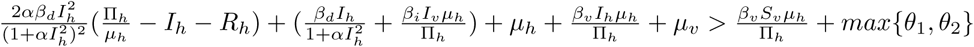 where,

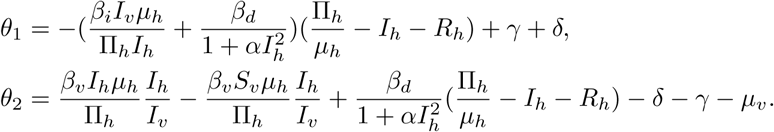

Then, the unique endemic equilibrium Y^*^ of the system (2.1) is globally asymptotically stable in region Θ.

#### Proof.

We show the global stability of *Y*^*\**^ using the Muldowney’s Theorem 4.2. We have to prove the following to apply the theorem:

1. There exists a compact absorbing set *K* in the interior of Θ.
2. There is a number *ν* > 0 such that the Lozinskii measure satisfies *µ*(*A*) < *-ν*.

The uniform persistence together with the boundedness of Θ is equivalent to the existence of an absorbing compact set in the interior of Θ(22). Now we have to prove that the Lozinskii measure satisfies *µ*(*A*) < *– ν* for *ν* > 0.

The jacobian matrix *J* of the system

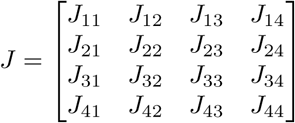

The definition of the second additive compound matrix is given by

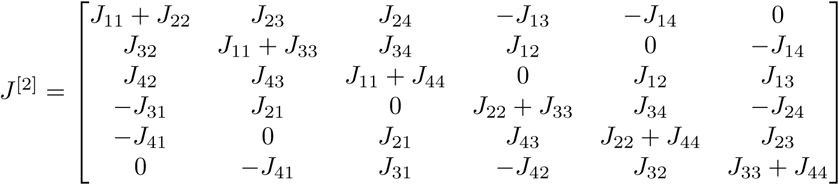

Therefore, the second additive compound matrix of *J* of our system is given by

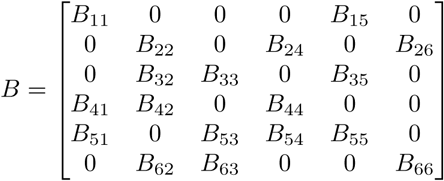

where

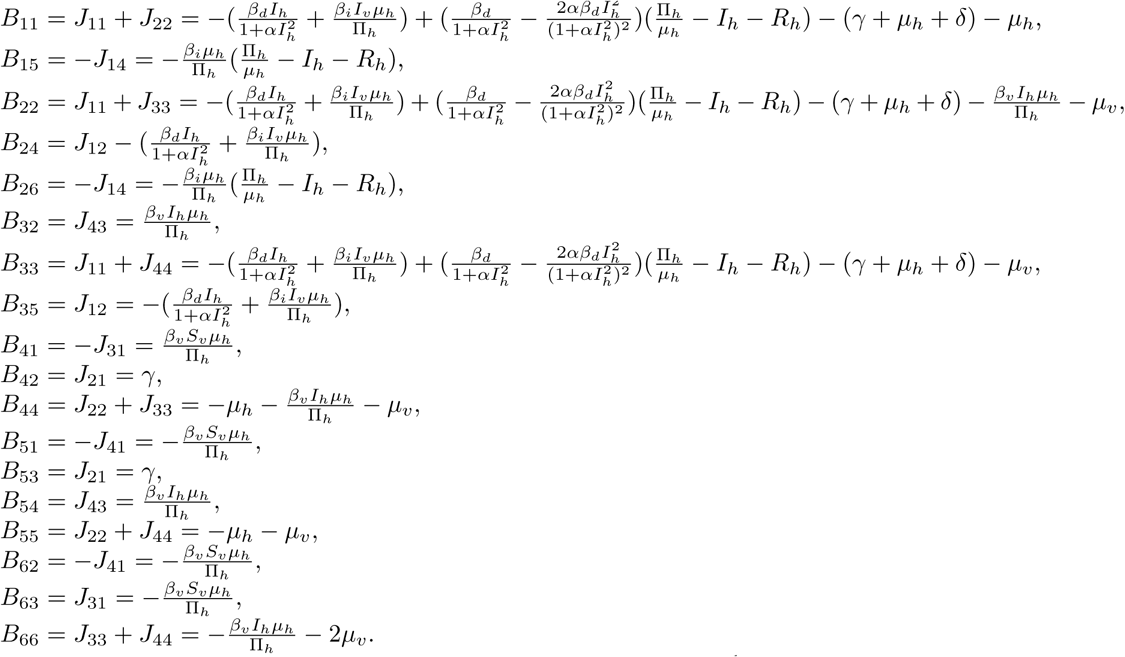

Let *Q*(*t*) be the following matrix of 6 *×* 6 which is invertible and *C*^1^

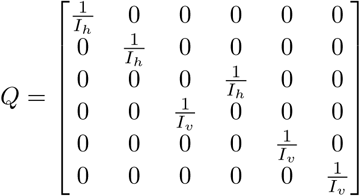

We have

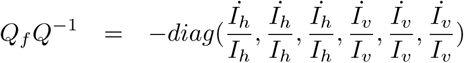

where *Q*_*f*_ is obtained by replacing each entry *Q*_*ij*_ of *Q* with the derivative of *Q*_*ij*_ in the direction of the vector field given by the system (2.1).

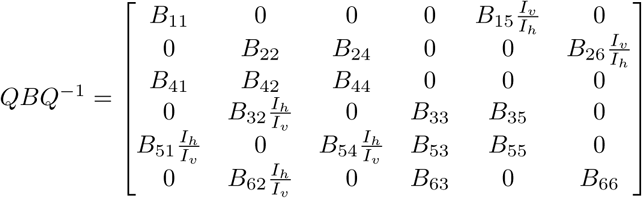

From the system, we have

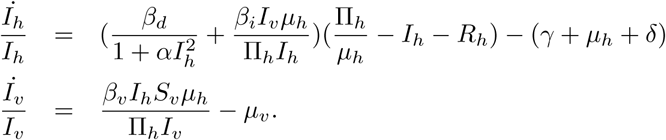

Then we obtain the matrix *A* = *Q*_*f*_ *Q*^*-*1^ + *QBQ*^*-*1^

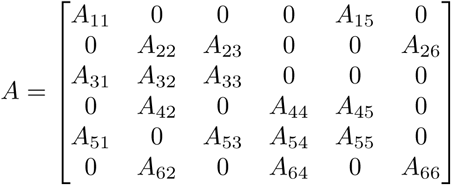

where,

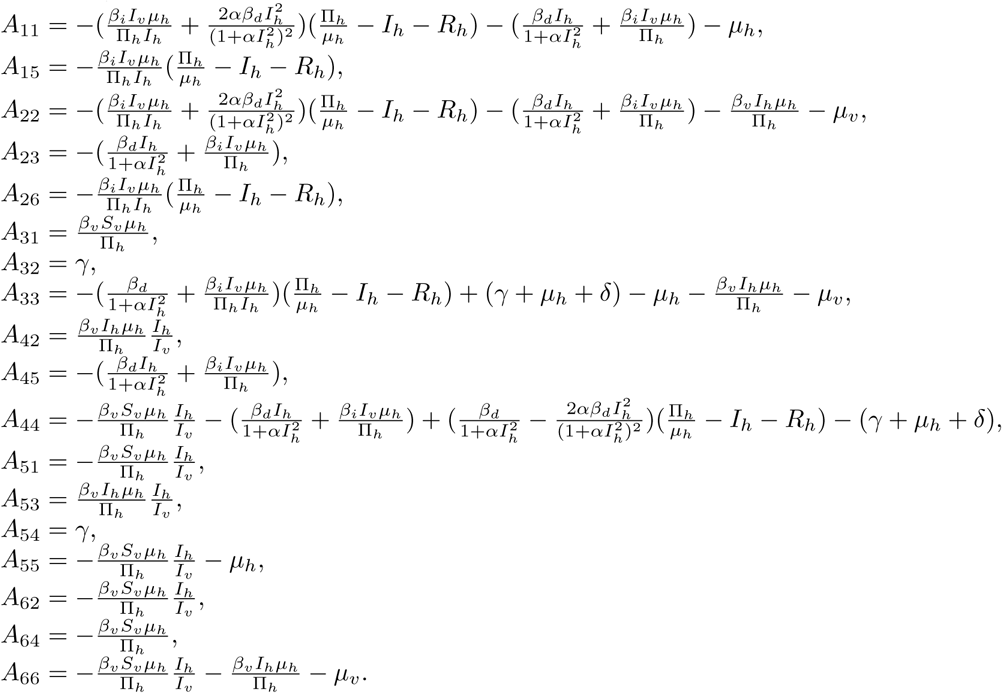

The second compound system of model (4.4) can demonstrated by the following system

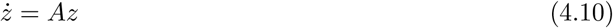

Let us define the norm on *R*^6^

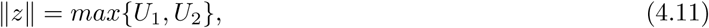

as in (23; 24) where *z* = (*z*_1_, *z*_2_, *z*_3_, *z*_4_, *z*_5_, *z*_6_) *∈ R*^6^, *U*_1_(*z*_1_, *z*_2_, *z*_3_) and *U*_2_(*z*_4_, *z*_5_, *z*_6_) are

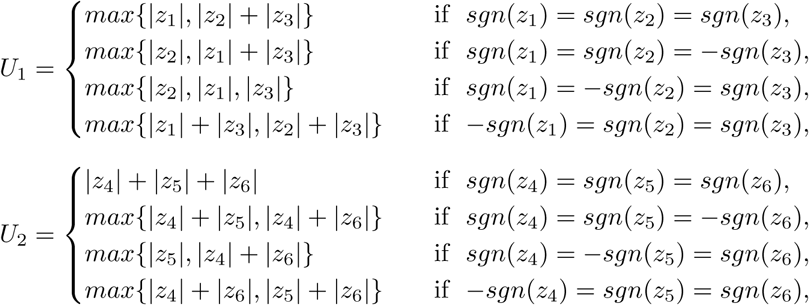

where the function *sgn*(*x*) is given by

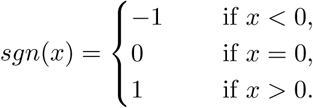

By definition of the norm, the following inequalities are obtained

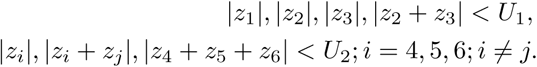

Following (25; 19), the Lozinskii measure for all solutions of *Ż* = *Az* can be evaluated as

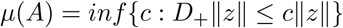

where *D*_+_ is the right hand derivative.

Now we have to evaluate *D*_+_ ||*z||*. To evaluate this quantity, sixteen different cases will arise according to the different octant and the definition of the norm (4.11) in each octant.

If *U*_1_ > *U*_2_, then nine cases will arise. In order to illustrate the evaluation procedures, we will provide detailed calculation of the first two cases.

#### Case 1

*z*_1_, *z*_2_, *z*_3_ > 0 and *|z*_1_*|* > *|z*_2_*|* + *|z*_3_*|* then

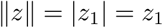

We have

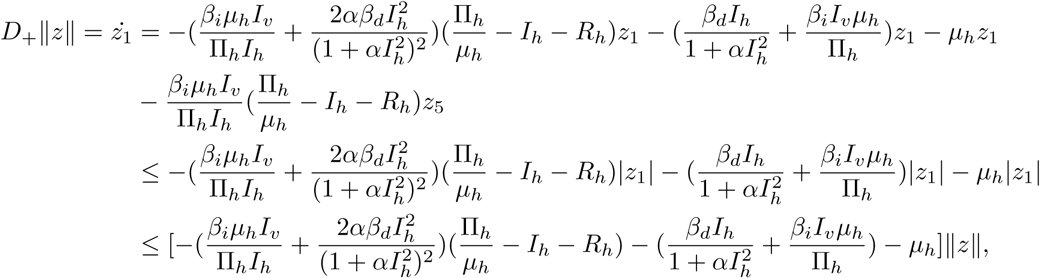

using that *|z*_5_*|* < *U*_2_ < *|z*_1_*|*.

#### Cases 2

*z*_1_, *z*_2_, *z*_3_ > 0 and *|z*_1_*|* < *|z*_2_*|* + *|z*_3_*|* then

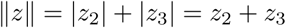

We have

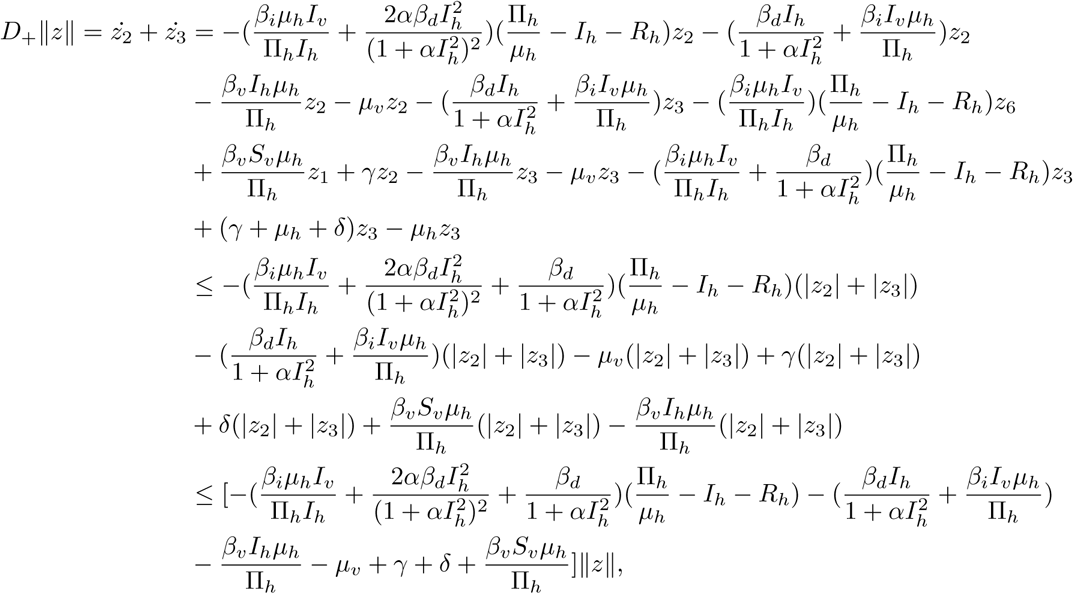

using that *|z*_6_*|* < *U*_2_ < *|z*_2_*|* + *|z*_3_*|* and *|z*_1_*|* < *|z*_2_*|* + *|z*_3_*|*.

#### Cases 3

*z*_1_ < 0, *z*_2_, *z*_3_ > 0 and *|z*_1_*|* > *|z*_2_*|* then

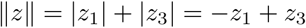

We have

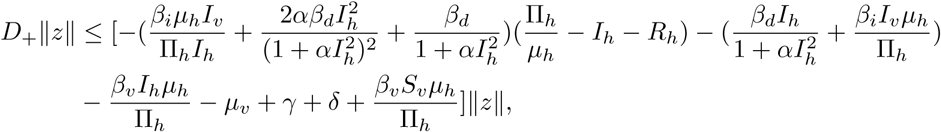

using that *|z*_5_*|* < *U*_2_ < *|z*_1_*|* + *|z*_3_*|*

#### Cases 4

*z*_1_ < 0, *z*_2_, *z*_3_ > 0 and *|z*_1_*|* < *|z*_2_*|* then

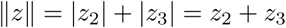

We have

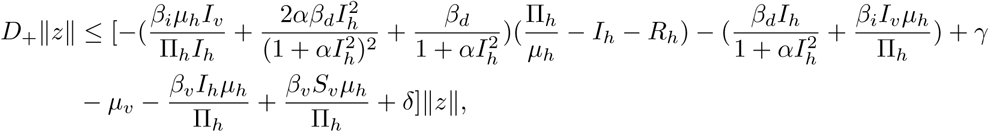

using that *|z*_6_*|* < *U*_2_ < *|z*_2_*|* + *|z*_3_*|*

#### Cases 5

*z*_1_, *z*_2_ > 0, *z*_3_ < 0 and *|z*_2_*|* > *|z*_1_*|* + *|z*_3_*|* then

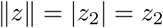

We have

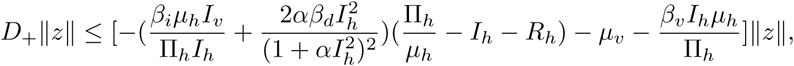

using that *|z*_6_*|* < *U*_2_ < *|z*_2_*|*

#### Cases 6

*z*_1_, *z*_2_ > 0, *z*_3_ < 0 and *|z*_2_*|* < *|z*_1_*|* + *|z*_3_*|* then

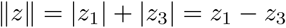

We have

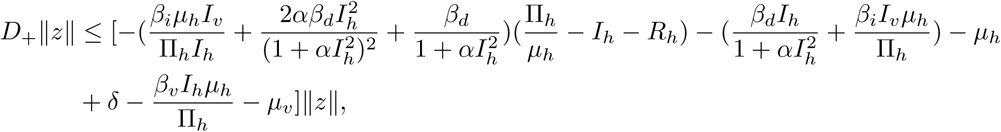

using that *|z*_5_*|* < *U*_2_ < *|z*_1_*|* + *|z*_3_*|*

#### Cases 7

*z*_1_, *z*_3_ > 0, *z*_2_ < 0 and *|z*_1_*|* > *max*{*|z*_2_*|, |z*_3_*|*} then

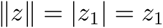

We have

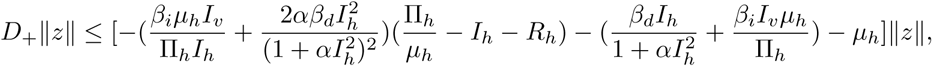

using that *|z*_5_*|* < *U*_2_ < *|z*_1_*|*

#### Cases 8

*z*_1_, *z*_3_ > 0, *z*_2_ < 0 and *|z*_2_*|* > *max*{*|z*_1_*|, |z*_3_*|*} then

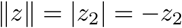

We have

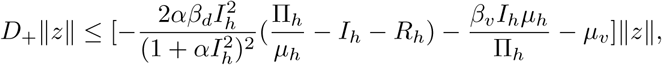

using that *|z*_6_*|* < *U*_2_ < *|z*_2_*|*

#### Cases 9

*z*_1_, *z*_3_ > 0, *z*_2_ < 0 and *|z*_3_*|* > *max*{*|z*_1_*|, |z*_2_*|*} then

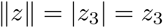

We have

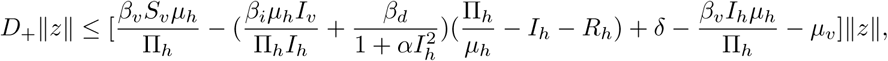

When *U*_1_ < *U*_2_, we have seven cases. We will do first two in detail.

#### Cases 10

*z*_4_, *z*_5_, *z*_6_ > 0 then

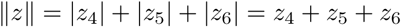

We have

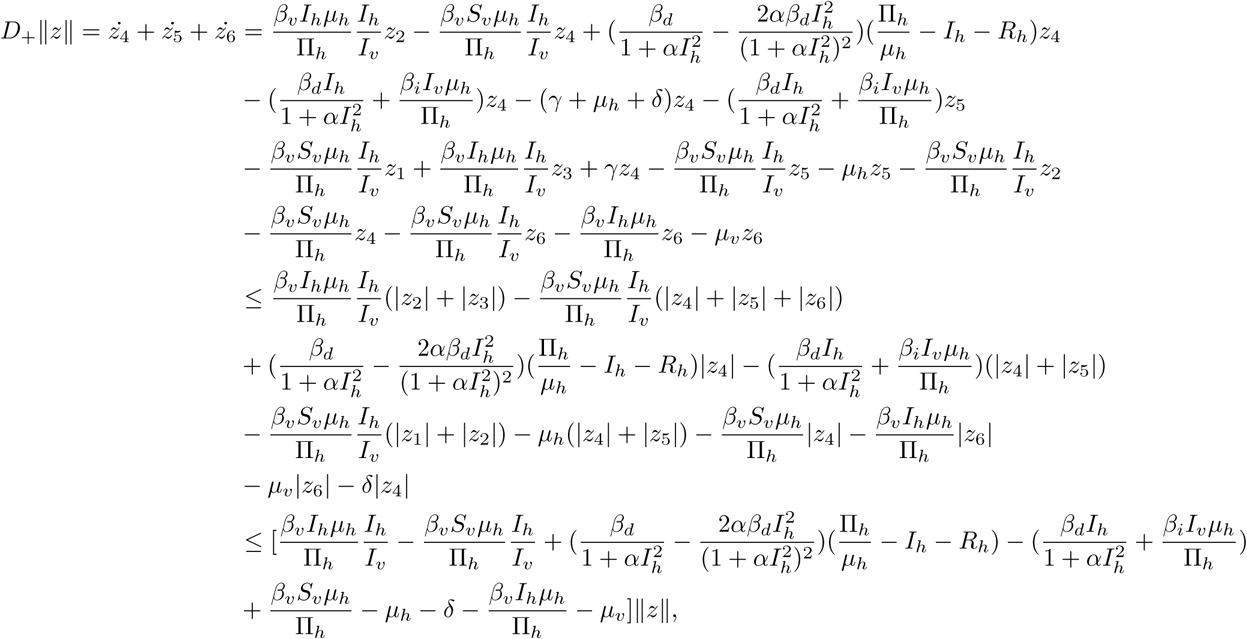

Using that *|z*_1_*|* < *U*_1_ < *|z*_4_*|*+*|z*_5_*|*+*|z*_6_*|, |z*_2_ +*z*_3_*|* < *U*_1_ < *|z*_4_*|*+*|z*_5_*|*+*|z*_6_*|* and *|z*_2_*|* < *U*_1_ < *|z*_4_*|*+*|z*_5_*|*+*|z*_6_*|*.

#### Cases 11

*z*_4_, *z*_5_ > 0, *z*_6_ < 0 and *|z*_5_*|* < *|z*_6_*|* then

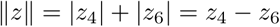

We have

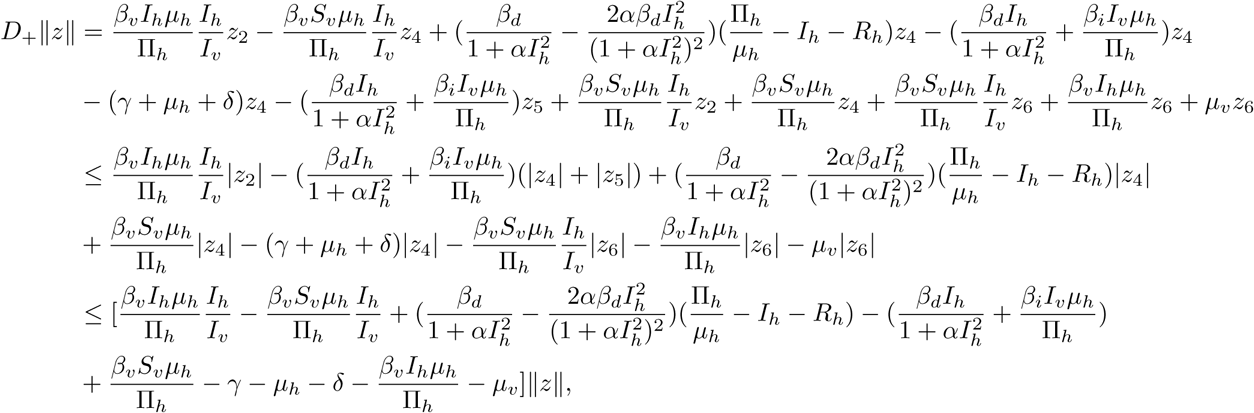

Using that *|z*_2_*|* < *U*_1_ < *|z*_4_*|* + *|z*_6_*|, |z*_4_*|, |z*_6_*|* < *|z*_4_*|* + *|z*_6_*|* and *|z*_4_ + *z*_5_*|* < *|z*_4_*|* + *|z*_6_*|*.

#### Cases 12

*z*_4_, *z*_5_ > 0, *z*_6_ < 0 and *|z*_5_*|* > *|z*_6_*|* then

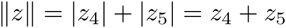

We have

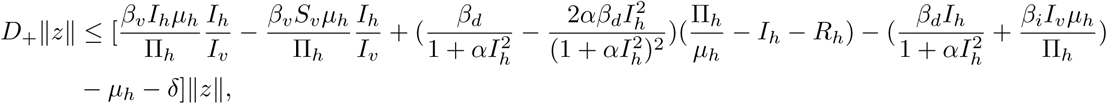

*|z*_2_ + *z*_3_*|* < *U*_1_ < *|z*_4_*|* + *|z*_5_*|* and *|z*_1_*|* < *U*_1_ < *|z*_4_*|* + *|z*_5_*|*.

#### Cases 13

*z*_4_, *z*_6_ > 0, *z*_5_ < 0 and *|z*_5_*|* > *|z*_4_*|* + *|z*_6_*|* then

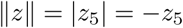

We have

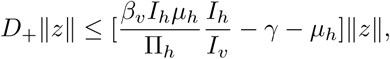

*|z*_1_*|* < *U*_1_ < *|z*_5_*|* and *|z*_3_*|* < *U*_1_ < *|z*_5_*|*.

#### Cases 14

*z*_4_, *z*_6_ > 0, *z*_5_ < 0 and *|z*_5_*|* < *|z*_4_*|* + *|z*_6_*|* then

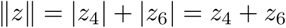

We have

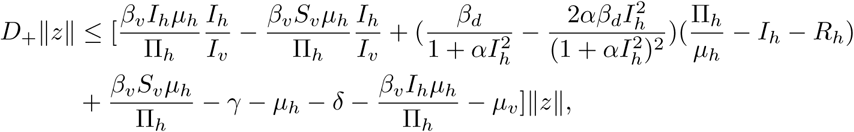

#### Cases 15

*z*_5_, *z*_6_ > 0, *z*_4_ < 0 and *|z*_5_*|* < *|z*_4_*|* then

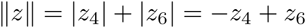

We have

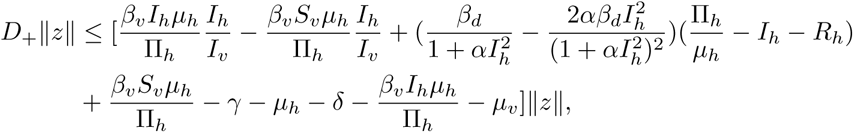

#### Cases 16

*z*_5_, *z*_6_ > 0, *z*_4_ < 0 and *|z*_5_*|* > *|z*_4_*|* then

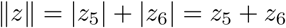

We have

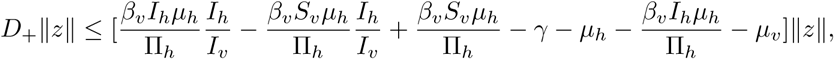

*|z*_1_| < *U*_1_ < |*z*_5_| + |*z*_6_| and |*z*_2_| < *U*_1_ < |*z*_5_| + |*z*_6_|.

From all cases. we obtain the following estimate

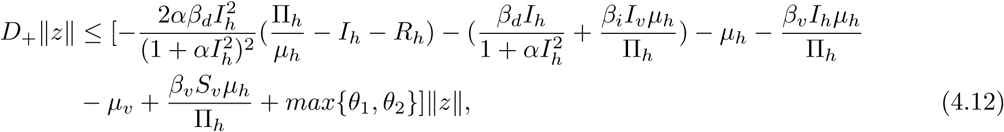

where,

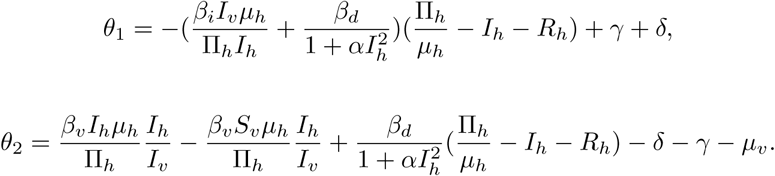

Since by the hypothesis

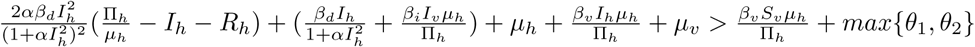

we concluded that

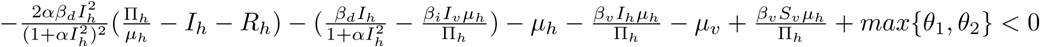

From (4.12), there exist *ν* > 0 such that

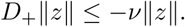

Therefore the Lozinskii measure satisfies

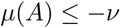

for all *x* = (*I*_*h*_, *R*_*h*_, *S*_*v*_, *I*_*v*_) *∈ K*. This completes the proof.

## 5. Bifurcation analysis

### 5.1. Backward bifurcation

There have been major implications of backward bifurcation in epidemiology. It has been well accepted that *R*_0_ < 1 is an essential requirement for eliminating the disease. But this opinion is recently challenged by some recent literature (26; 27; 28). The occurrence of backward bifurcation phenomenon indicates that the disease may persist even if this condition is satisfied. Alternatively, this phenomenon shows that if *R*_0_ < 1, although the DFE is stable, another stable endemic equilibrium may coexist simultaneously. When the multiple stable equilibrium coexist then the population will reach the final equilibrium depending on the initial conditions. Backward bifurcation phenomenon for the system (2.1) is shown in Fig. 1. Here, a stable DFE and a stable endemic equilibrium point coexist when 0 < *R*_*c*_ < *R*_0_ < 1, where *R*_*c*_ is the critical value, which is shown in Fig. 1 and *R*_*c*_ = 0.274 for the given parameters values. In Fig. 1, the green line represents the unstable equilibrium, while the blue line represents the stable equilibrium. The disease free equilibrium is stable when *R*_0_ < *R*_*c*_. This phenomenon is very interesting when *R*_0_ changes slowly. For instance, when the infected host population is very close to extinction 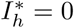 i.e for DFE, then *R*_0_ increases slowly. But when *R*_0_ crosses the threshold quantity *R*_0_ = 1, the number of infected host population will jump to large endemic equilibrium. Again when the infected population is very close to the endemic equilibrium, *R*_0_ decreases slowly through the threshold point *R*_0_ = 1, rather than switching to the diseases free equilibrium, the infective population remain close and converge to the stable endemic equilibrium. As soon as *R*_0_ decreases below the threshold *R*_*c*_, the number of infected population immediately jumps towards zero to diseases free equilibrium, while the endemic equilibrium disappear.

**Figure 1:**
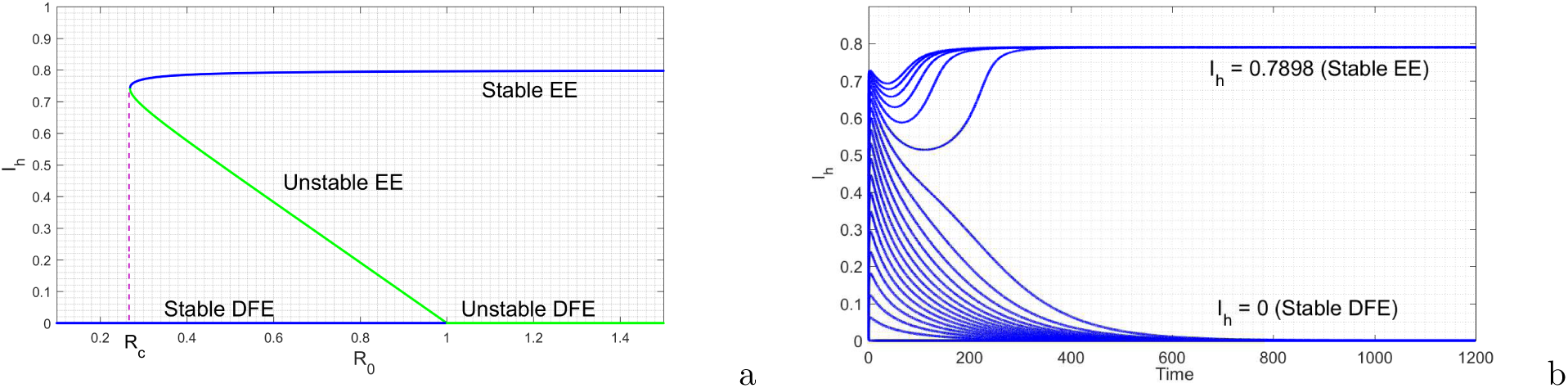
Backward bifurcation diagram for infected host at *R*_0_ = 1 where the parameter values are given by: Π_*h*_ = 6, *α* = 0.66, *β*_*d*_ = 0.01, *β*_*v*_ = 0.00072, *µ*_*h*_ = .5, *µ*_*v*_ = 0.02, Π_*v*_ = 50, *δ* = 7, *γ* = 0.0004. Time series plot using different initial conditions of the model. Parameter values are same as backward bifurcation diagram except *β*_*i*_ = 0.5.

Following Castillo-Chavez and Song (27), the phenomenon of backward bifurcation of the system(2.1) is established analytically. The Jacobian of the system (3.2) at diseases free equilibrium *Y*_0_ is given as

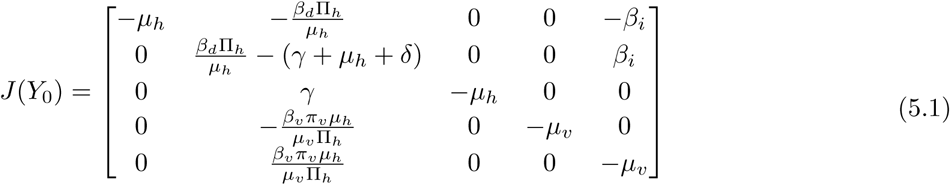

Let *β*_*i*_ be the bifurcation parameter and using *R*_0_ = 1, we have

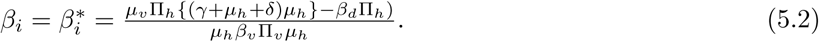

Here 0 is the simple eigen value and the jacobian *J* (*Y*_0_) at 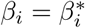 has a right eigen vector corresponding to zero eigen value is given by *w* = (*w*_1_, *w*_2_, *w*_3_, *w*_4_, *w*_5_) and has a left eigen vector corresponding to zero eigen vector is given by *v* = (*v*_1_, *v*_2_, *v*_3_, *v*_4_, *v*_5_), where

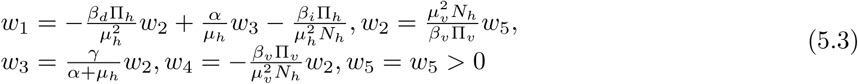

and

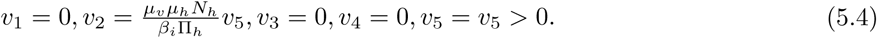

The following second order partial derivatives of *g*_*i*_ at DFE *Y*_0_ are calculated as follows:

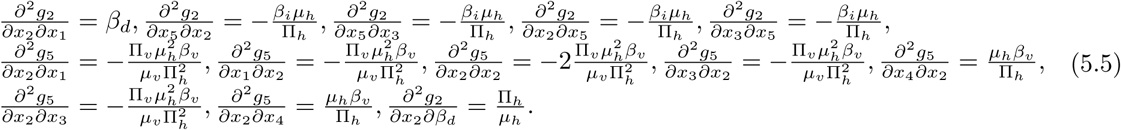

Now the coefficients *a* and *b* defined in the theorem of Castillo-chavez and Song are calculated as follows

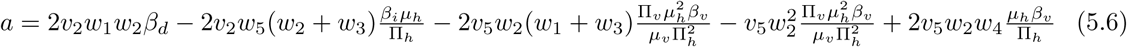

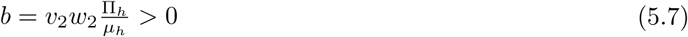

We see that the coefficient *b* is always positive, hence the system (2.1) undergoes backward bifurcation at *R*_0_ = 1, if *a* > 0.

The backward bifurcation phenomenon would occur for those values of *R*_0_ such that *R*_*c*_ < *R*_0_ < 1. This is illustrated by numerical simulation with the following set of hypothetical parameter values: Π_*h*_ = 6, *α* = 0.66, *β*_*d*_ = 0.01, *β*_*v*_ = 0.00072, *β*_*i*_ = 0.5, *µ*_*h*_ = .5, *µ*_*v*_ = 0.02, Π_*v*_ = 50, *δ* = 7, *γ* = 0.0004. With the set of this parameters, *R*_*c*_ = 0.274 and *R*_0_ = 0.5160 < 1 (so that *R*_*c*_ < *R*_0_ < 1). Corresponding bifurcation diagram is plotted in the Fig 1(a). A time series of 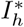 is also plotted in the Fig 1(b), showing the DFE (corresponding to 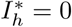) and two endemic equilibria. Further, the Fig 1(a) shows that one of the endemic equilibrium 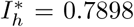 is locally asymptotically stable, and the other is unstable, and the DFE is is locally asymptotically stable. This phenomenon clearly shows that the co-existence of two locally asymptotically stable equilibrium when *R*_0_ < 1. This confirm that the model (2.1) undergoes the backward bifurcation phenomenon.

### 5.2. Hysteresis

For forward transcritical bifurcation usually a model has two locally stable branches, one is DFE when *R*_0_ < 1, which is locally asymptotically stable and another is endemic equilibrium when *R*_0_ > 1, which is stable. But there may be more than one stable endemic equilibrium for the model when *R*_0_ > 1. It is possible that more than one endemic equilibrium coexist in epidemic models even when *R*_0_ > 1. This leads to an unusual phenomenon of forward bifurcation with hysteresis effect, which is shown numerically for our model (2.1). We find that our model exhibits a hysteresis effect where multiple endemic equilibrium coexist for *R*_0_ > 1 (see Fig. 2). The blue line represents two outer equilibrium, which is stable while the green line represent the interior equilibrium, which is unstable.

**Figure 2:**
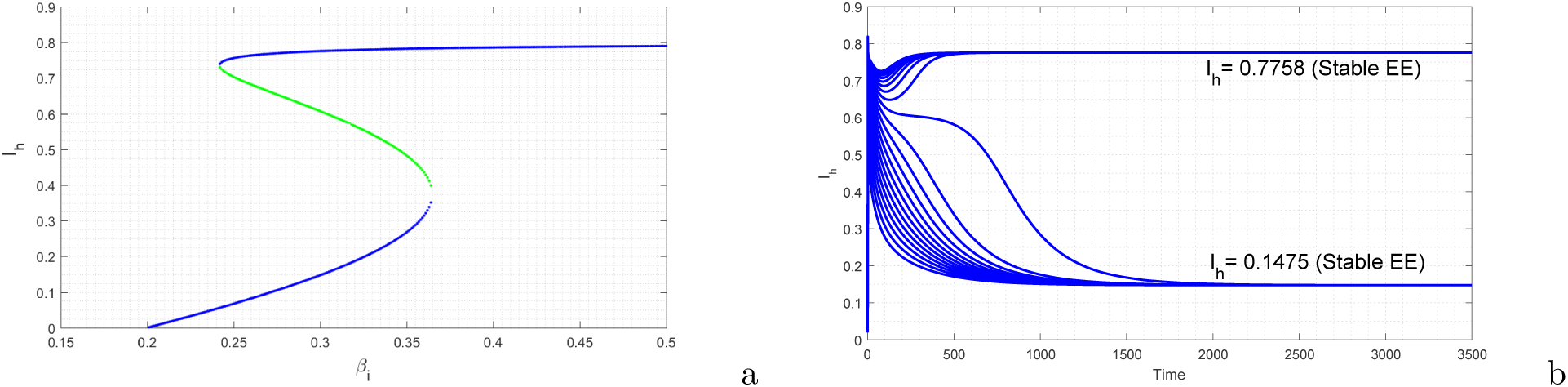
Hysteresis diagram of the model. Parameters are given by: Π_*h*_ = 6, *α* = 0.66, *β*_*d*_ = .5, *β*_*v*_ = 0.00072, *µ*_*h*_ = .5, *µv* = 0.02, Π*v* = 50, *δ* = 7, *γ* = 0.0004. Time series plot using different initial conditions of the model. Parameters are same as hysteresis except *β*_*i*_ = 0.3.

Further investigation reveals that the endemic equilibrium may also be stable in a region where *R*_0_ < 1 (see Fig. 3). This indicates that the disease may persist for *R*_0_ < 1, even if the type of transcritical bifurcation at *R*_0_ = 1 is forward.

**Figure 3:**
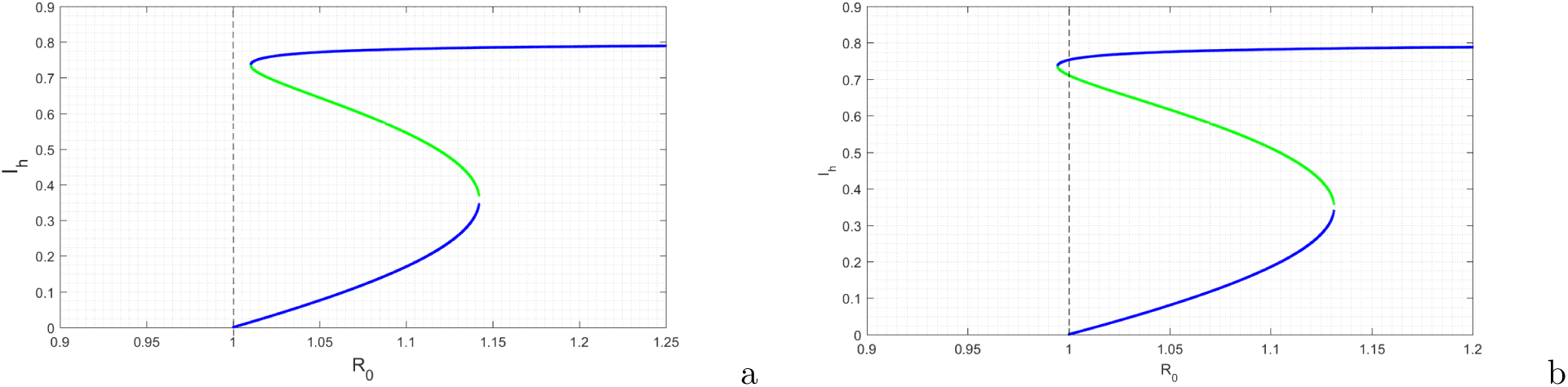
Hysteresis diagram of the model. Parameter values are given by: Π_*h*_ = 6, *α* = 0.66, *β*_*v*_ = 0.00072, *µ*_*h*_ = .5, *µ*_*v*_ = 0.02, Π_*v*_ = 50, *δ* = 7, *γ* = 0.0004, *β*_*i*_ = 0.3, *β*_*d*_ = 0.48(*for*(*a*)), *β*_*d*_ = 0.47 (*for*(*b*)).

## 6. Numerical scenarios

### 6.1. Impact of temporary control on bistable dynamics

The model (2.1) undergoes forward bifurcation with hysteresis as shown in Fig. 2. This phenomenon is rarely observed in disease models and therefore this finding will definitely help getting more insight into the transmission patterns. From Fig. 2, it is observed that the system approaches two different levels of endemicity depending on the initial conditions. Thus, the lower and upper equilibria have disjoint basin of attractions. In this situation, temporary control measures may be implemented to keep the equilibrium level of infected population to low endemicity. For backward bifurcation case, we can employ temporary control interventions to push the solution into the basin of attraction of the DFE. Similarly, in case of hysteresis, these control measures will compel the solutions to the basin of attraction of the lower endemic equilibrium. Here, we control the indirect transmission rate (*β*_*i*_) for a certain period of time, to force the solutions of the system to lower endemic value. We use the following to model this

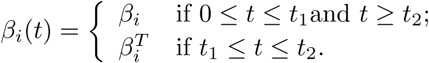

For backward bifurcation case, we observe that a solution of the system approaches the upper equilibrium whenever no control interventions are applied. Further, the solution experience a sudden jump to the upper equilibrium after initial decline for relatively short-term control measures. However, with the same initial conditions the equilibrium shifts down to the lower equilibrium, if the duration and/or strength of the control measure is large enough (see Fig. 4(a)). Similarly for the hysteresis case suitable strength of control measure can push the higher endemic level of infectives to lower equilibrium (see Fig. 4(b)). These results suggest that, suitable application of control will thus push the solution to the basin of attraction of lower equilibrium or DFE. Therefore, temporary control strategies will be beneficial in the bistable region of the solution space. Furthermore, delay in applying temporary control will have crucial effect on the behavior of the system. Early temporary control will drive the solution to the lower equilibrium whereas delayed temporary control with same strength and duration will keep the solution in the basin of attraction of the upper equilibrium.

**Figure 4:**
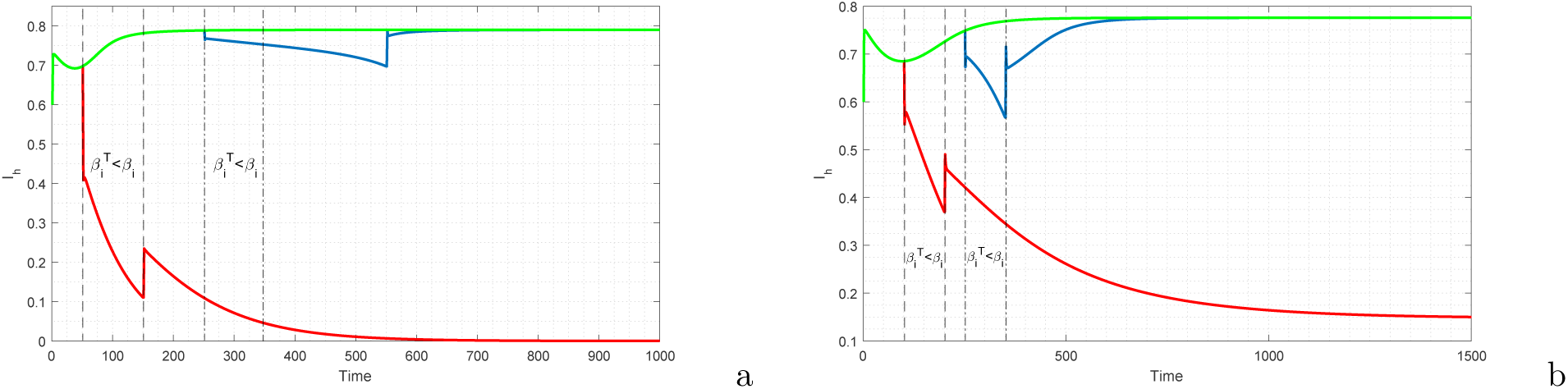
Impact of temporary control on bi-stable dynamics.

### 6.2. Impact of varying the strength of psychological effect (α)

We investigate the effect of the parameter *α* which describes the behavioral change of the susceptible hosts towards the infected hosts. It is observed that the basic reproduction number does not depend explicitly on *α*. But, numerical simulations reveal that when the disease is endemic, the steady state value of the infected hosts decrease as *α* increases (as shown in Fig. 5).

**Figure 5:**
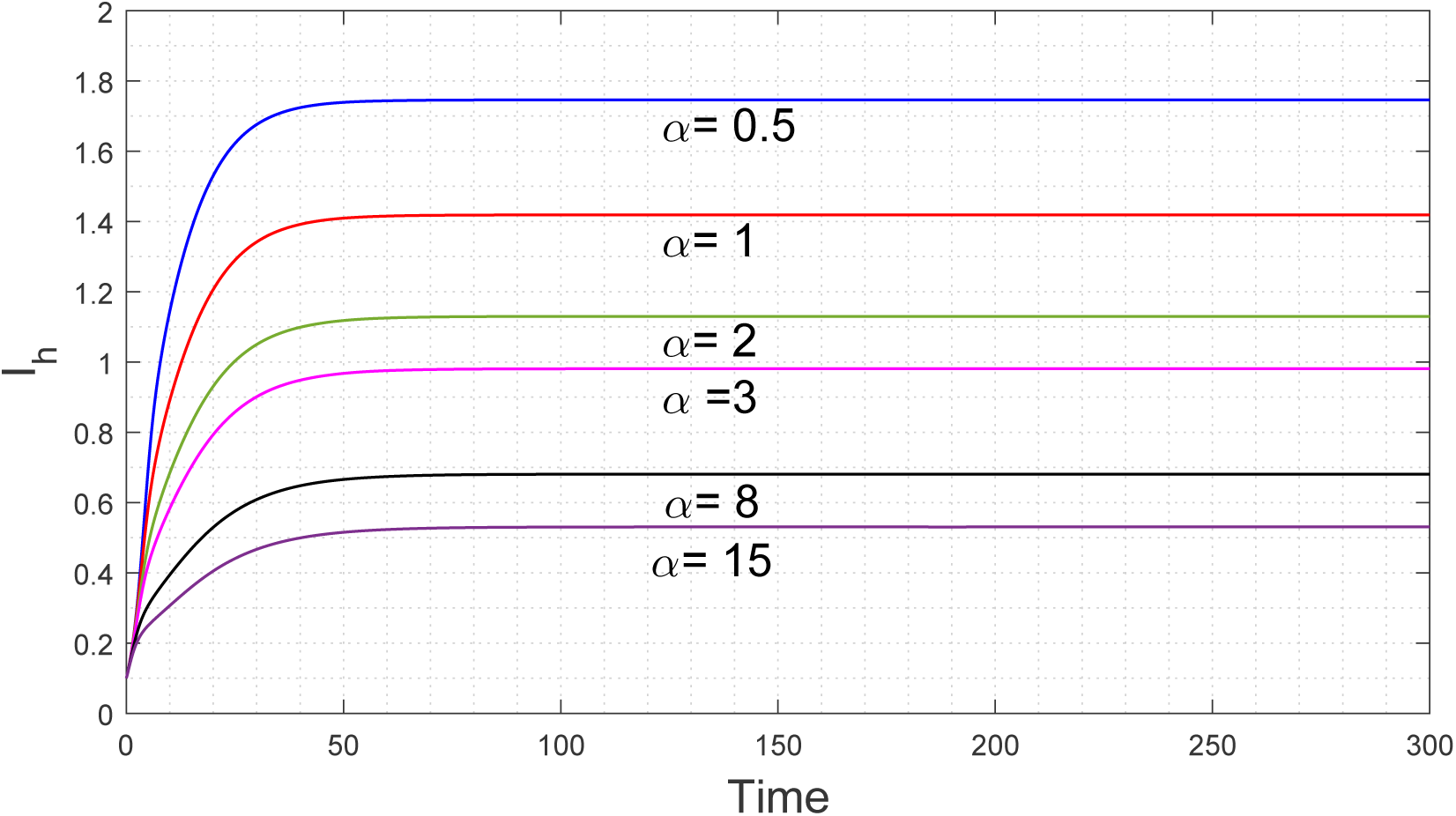
The dependence of *I*^*\**^ on the parameter *α*

Further, In Fig. 6, we show the variation in the concentration of infected host with respect to *α* (psychological parameters)and *β*_*d*_ (direct transmission rate). It is apparent from this figure that with an increase in the values of *β*_*d*_, the concentration of infected host increases. On the other hand, with an increase of the values of *α*, the concentration of infected host decreases.

**Figure 6:**
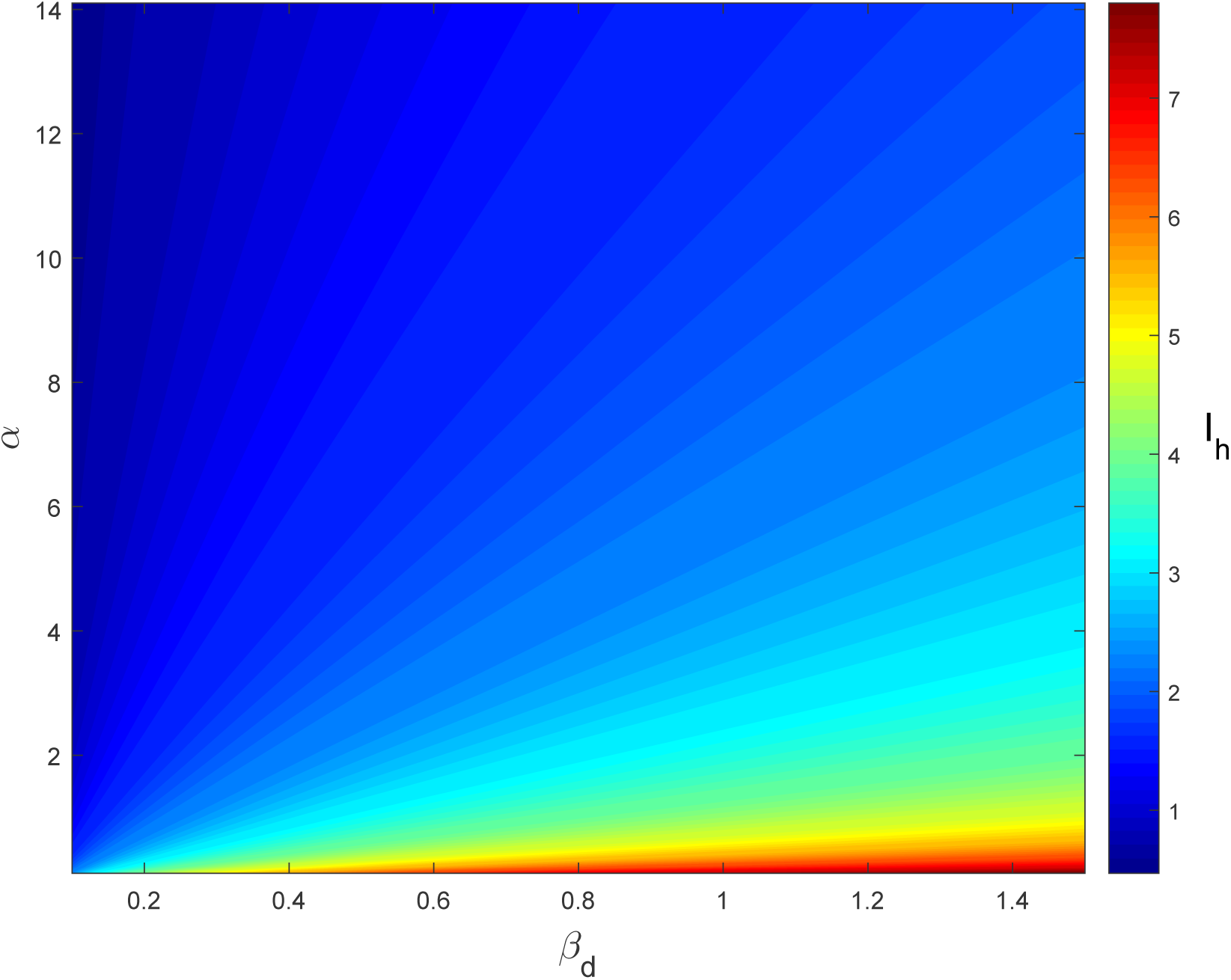
The contour plot of infected host in terms of the parameters: *β*_*d*_ (infection rate for infected host to susceptible host population) and *α* (psychological parameter)

These results suggest that *α* has a significant effect on decreasing the endemic equilibrium state of the infected host population.

### 6.3. Effect of heterogeneous indirect transmission

The vector-host interaction is assumed to be well-mixed in the proposed model 2.1, but this assumption is violated when we consider a sufficiently large spatial scales. It is evident that in a large area, a given vector may only have the opportunity to contact a small subset of hosts (29). To incorporate heterogeneity in our model, we adopt the method followed by Kong et. al (30). Thus, the transmission term from infectious vector to susceptible hosts is given by:

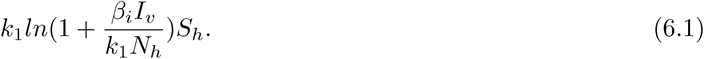

Similarly, the transmission term from infectious hosts to susceptible vectors is given by:

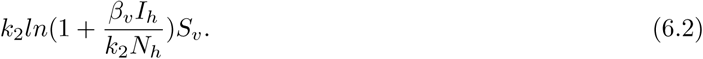

Here *k*_1_ and *k*_2_ are the levels of heterogeneity of vector-host and host-vector transmissions respectively. Since *k*_1_ and *k*_2_ both are the levels of heterogeneity, we assumed that *k*_1_ = *k*_2_ = *k* for simplicity. Therefore, replacing these heterogeneity terms into our model 2.1, we obtain the following system of ordinary differential equations:

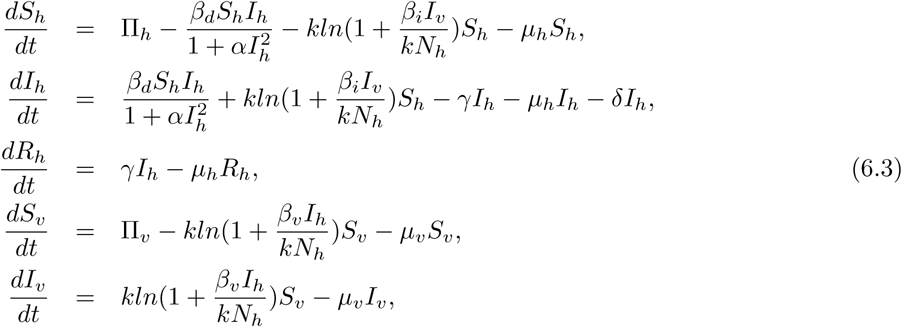

Using numerical simulations, the impact of the heterogeneity parameter *k* have been studied. We computed the time-series of infected hosts for different values of *k*: 0.01, 0.06, 0.11, 0.16. Keeping the initial conditions fixed, simulations are shown in Fig. 7. The corresponding homogeneous mixing model solution is also plotted to compare it with different levels of heterogeneity.

**Figure 7:**
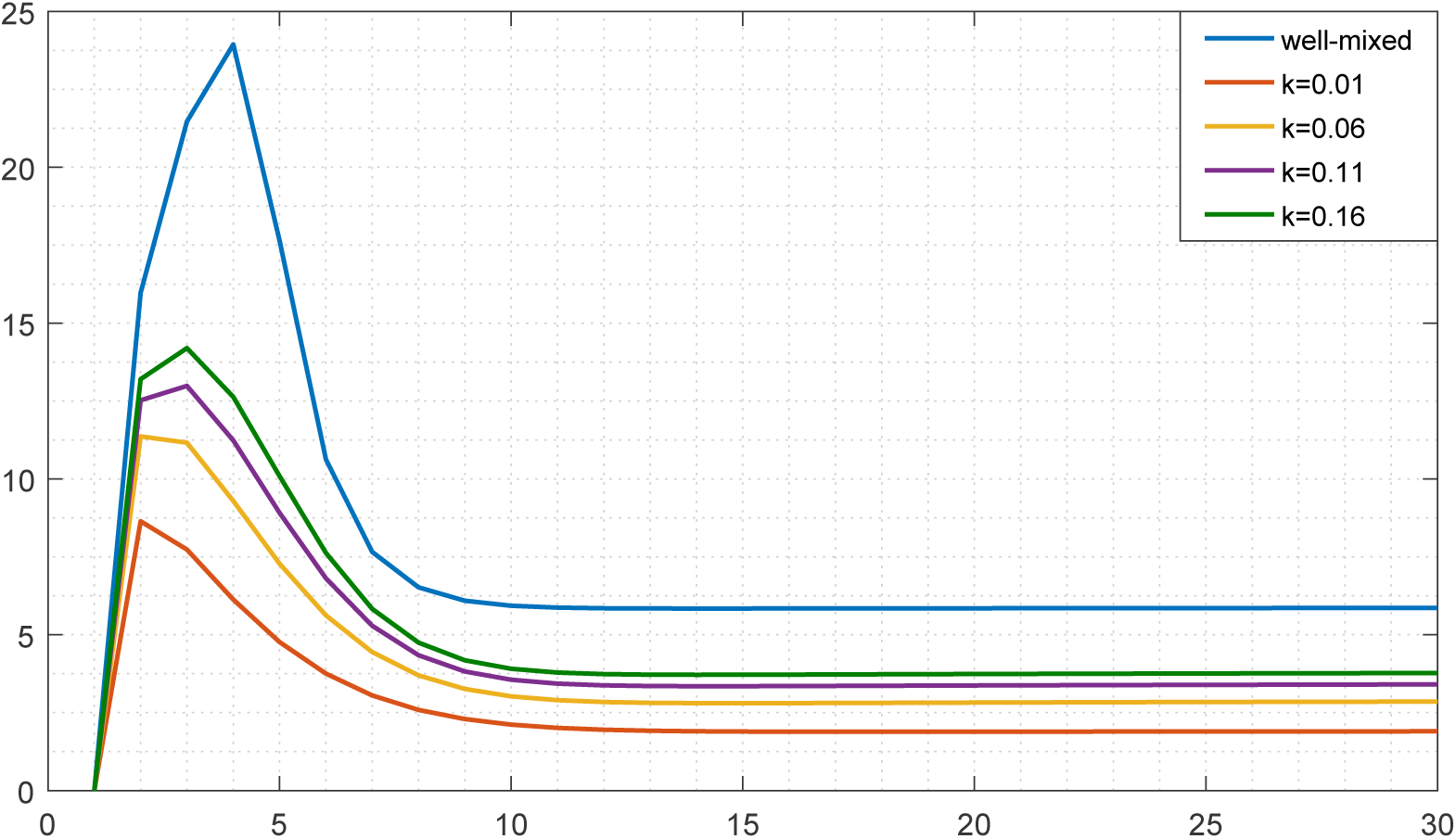
Time series of infected hosts for different values of *k*.

From Fig. 7, it is observed that the peak infected host levels are decreased with increase in the heterogeneity levels. Epidemiologically, lower values of *k* indicates that a relatively low proportion of susceptibles has the chance to contact vectors. For this the peak infected hosts decreases with lower values of *k*. Thus, high level of heterogeneity increases the extinction probability of the disease.

Further, we investigated the sensitivity of the parameters *k, β*_*i*_ and *β*_*v*_ with respect to the equilibrium susceptible and infected host levels, Table 4. It is observed that the equilibrium infected host level increases with increase in the parameter *k*.

**Table 4:** Variation in the equilibrium susceptible host level and equilibrium infected host level due to changes in the parameters *k, β*_*i*_ and *β*_*v*_.

| $k$ | $\beta_i$ | $\beta_v$ | Equilibrium susceptible host level | Equilibrium infected host level |
| --- | --- | --- | --- | --- |
| $k = 0.01$ | 0.9 | 0.0 | 8.8339 | 1.5767 |
|  | 0.0 | 0.82 | 8.8348 | 1.5763 |
|  | 0.9 | 0.82 | 8.1183 | 1.9331 |
| $k = 0.05$ | 0.9 | 0.0 | 8.8339 | 1.5767 |
|  | 0.0 | 0.82 | 8.8348 | 1.5763 |
|  | 0.9 | 0.82 | 6.4094 | 2.7842 |
| $k = 0.1$ | 0.9 | 0.0 | 8.8339 | 1.5767 |
|  | 0.0 | 0.82 | 8.8348 | 1.5763 |
|  | 0.9 | 0.82 | 5.1622 | 3.4053 |
| $k = 100$ | 0.9 | 0.0 | 8.8339 | 1.5767 |
|  | 0.0 | 0.82 | 8.8348 | 1.5763 |
|  | 0.9 | 0.82 | 0.1922 | 5.8804 |

## 7. Discussion

In this study, by combining stability and bifurcation analysis we have examined the dynamical properties of an epidemic model with two routes of transmission. The basic reproduction number is derived for the proposed model. It is shown that the local dynamics of the system is governed by the threshold quantity *R*_0_. Local and global stability conditions of the endemic equilibrium are derived. Following a standard theorem of Castillo-Chavez and Song (27), we established the occurrence of backward bifurcation for the proposed model. Further, we verified the phenomena of backward bifurcation and forward bifurcation with hysteresis numerically. Furthermore, under some certain conditions it is found that the epidemic growth rate is more sensitive to direct transmission (*β*_*d*_) than to indirect transmission. This result is then verified using normalized sensitivity index. Finally, we investigate the impact of temporary control on the bistable behaviour of the system. Numerical experiments reveal that it is indeed possible to switch the stability of equilibria. In case of backward bifurcation, temporary control can curb the disease endemic equilibrium to disease free state. On the other hand, the system can be brought to lower endemic levels by the employment of temporary control. We found that the strength of psychological effect will have significant contribution in lowering the endemic infection level of the host population. Further investigation reveals that increase in transmission heterogeneity will ensure fast eradication of the disease.

In summary, global stability of the unique endemic equilibrium indicate that the disease will persist under certain parametrization of the system. Additionally, the diseases with two transmission routes can exhibit bi-stable dynamics. These bi-stable feature of the diseases may impose significant difficulties on disease control strategies. Moreover, efficient application of temporary control will be helpful in bi-stable regions. While investigating the effect of heterogeneity, we observe that high levels of heterogeneities will result in slower disease spread.

## Acknowledgement

The research work of Sk Shahid Nadim is supported by CSIR, Government of India, New Delhi in the form of senior research fellowship. Indrajit Ghosh was supported by the Senior Research Fellowship from University Grants Commission, Government of India. The funders had no role in study design, data collection and analysis, decision to publish or preparation of the manuscript.

## References

[1] G. Dubas, L.M. Carvalho, A. Rmabaut, T. Bedford, MERS-CoV spillover at the camel-human interface, ELife 7 (2018) e31257.

[2] D. Gao, Y. Lou, D. He, T.C. Porco, Y. Kuang, G. Chowell, S. Ruan, Prevention and control of Zika as a mosquito-borne and sexually transmitted disease: a mathematical modeling analysis, Scientic reports 6 (2016) 28070.

[3] G.M. Leunng, T.H. Lam, L.M. Ho, S.Y. Ho, B.H.Y. Chan, I.O.L. Wong, A.J. Hedley, The impact of community psychological responses on outbreak control for severe acute respiratory syndrome in Hong Kong, Journal of Epidemiology & Community Health 57(11) (2003) 857–863.

[4] F. Verelst, L. Willem, P. Beutels, Behavioural change models for infectious disease transmission: a systematic review (2010–2015), Journal of The Royal Society Interface 13(125) (2016) 20160820

[5] A.B. Gumel, S. Ruan, T. Day, J. Watmough, F. Brauer, P.V.D. Driessche, D. Gabrielson, C. Bowman, M.E. Alexander, S. Ardal, et al., Modelling strategies for controlling SARS outbreaks, Proceedings of the Royal Society of London B: Biological Sciences 271(1554) (2004) 2223–2232.

[6] W. Wang, S. Ruan, Simulating the SARS outbreak in Beijing with limited data, Journal of theoretical biology 227(3) (2004) 369–379.

[7] W. Liu, S.A. Levin, Y. Iwasa, Influence of nonlinear incidence rates upon the behavior of SIRS epidemiological models, Journal of mathematical biology 23(2) (1986) 187–204.

[8] S. Ruan, W. Wang, Dynamical behavior of an epidemic model with a nonlinear incidence rate, Journal of Differential Equations 188(1) (2003) 135–163.

[9] D. Xiao, S. Ruan, Global analysis of an epidemic model with nonmonotone incidence rate, Mathematical biosciences 208(2) (2007) 419–429.

[10] V.T. Covello, R.G. Peters, J.G. Wojtecki, R.C. Hyde, Risk communication, the West Nile virus epidemic, and bioterrorism: responding to the commnication challenges posed by the intentional or unintentional release of a pathogen in an urban setting, Journal of Urban Health 78(2) (2001) 382–391.

[11] V. Capasso, G. Serio, A generalization of the Kermack-McKendrick deterministic epidemic model, Mathematical Biosciences 42(1-2) (1978) 43–61.

[12] H.R. Thieme, Mathematics in population biology, Princeton University Press (2003).

[13] L. Cai, J. Xiang, X. Li, A.A. Lashari, A two-strain epidemic model with mutant strain and vaccination, Journal of Applied Mathematics and Computing 40(1-2) (2012) 125–142.

[14] J.H. Tien, D.J.D. Earn, Multiple transmission pathways and disease dynamics in a waterborne pathogen model, Bulletin of mathematical biology 72(6) (2010) 1506–1533.

[15] P.V.D. Driessche, J. Watmough, Reproduction numbers and sub-threshold endemic equilibria for compartmental models of disease transmission, Mathematical biosciences 180(1-2) (2002) 29–48.

[16] S.T.R.De. Pinho, C.P. Ferreira, L. Esteva, F.R. Barreto, V.C.M.e. Silva, M.G.L. Teixeira, Modelling the dynamics of dengue real epidemics, Philosophical Transactions of the Royal Society of London A: Mathematical, Physical and Engineering Sciences 368(1933) (2010) 5679–5693.

[17] Y. Li, S. James, On Bendixsons criterion, Journal of Differential Equations 106 (1993) 27–39.

[18] M.Y. Li, J.S. Muldowney, On RA Smith’s autonomous convergence theorem, The Rocky Mountain Journal of Mathematics (1995) 365–379.

[19] M.Y. Li, J.S. Muldowney, A geometric approach to global-stability problems, SIAM Journal on Mathematical Analysis 27(4) (1996) 1070–1083.

[20] G. Li, W. Wang, Z. Jin, Global stability of an SEIR epidemic model with constant immigration, Chaos, Solitons & Fractals 30(4) (2006) 1012–1019.

[21] H.I. Freedman, S. Ruan, M. Tang, Uniform persistence and flows near a closed positively invariant set, Journal of Dynamics and Differential Equations 6(4) (1994) 583–600.

[22] V. Huston, K. Schmitt, Permanence and the dynamics of biological systems, Mathematical biosciences 111(1) (1992) 1–71.

[23] B. Buonomo, C. Vargas-De-León, Global stability for an HIV-1 infection model including an eclipse stage of infected cells, Journal of Mathematical Analysis and Applications 385(2) (2012) 709–720.

[24] A.B. Gumel, C.C. McCluskey, J. Watmough, An SVEIR model for assessing potential impact of an imperfect anti-SARS vaccine (2006)

[25] R.H. Martin Jr, Logarithmic norms and projections applied to linear differential systems, Journal of Mathematical Analysis and Applications 45(2) (1974) 432–454.

[26] J. Arino, C.C. McCluskey, P.V.D. Driessche, Global results for an epidemic model with vaccination that exhibits backward bifurcation, SIAM Journal on Applied Mathematics 64(1) (2003) 260–276.

[27] C. Castillo-Chavez, B. Song, Dynamical models of tuberculosis and their applications, Mathematical biosciences and engineering 1(2) (2004) 361–404.

[28] I. Ghosh, T. Sardar, J. Chattopadhyay, A mathematical study to control visceral leishmaniasis: an application to South Sudan, Bulletin of mathematical biology 79(5) (2017) 1100–1134.

[29] N.D. Barlow, Non-linear transmission and simple models for bovine tuberculosis, Journal of Animal Ecology 69(4) (2000) 703–713.

[30] L. Kong, J. Wang, Z. Li, S. Li, Q. Liu, H. Wu, W. Yang, Modeling the heterogeneity of Dengue transmission in a city, International journal of environmental research and public health 15(6) (2018).

